# A bulk cell heterozygous knock-in strategy for targeted protein degradation

**DOI:** 10.64898/2026.05.19.726384

**Authors:** Beibei Liu, Chao Qi, Tomoharu Kanie

## Abstract

Targeted protein degradation using conditional degron tag (CDT) technology is a powerful method for rapidly degrading a protein of interest (POI) upon the addition of a degrader drug. A prerequisite for the temporally controlled degradation of an endogenous POI is the generation of homozygous knock-in cells with the degron tag integrated at either the N- or C-terminus of their gene loci. However, obtaining those homozygous knock-in cells often requires selecting many single-cell clones, as human cells typically exhibit low homology-directed repair (HDR) activities. Additionally, tagging a degron to an endogenous protein may inadvertently reduce protein expression, potentially affecting protein function even before the drug is administered. Here, we develop a method for generating degron-tagged knock-in cells that allows us to skip the laborious single-cell cloning. This method arose from our observation that most knock-in cells carry the degron tag only in one allele (heterozygous), while the other allele typically harbors a frameshift insertion/deletion. This observation allowed us to bypass the need for single-cell cloning. We validated our method by knocking in degron tags at the N-terminus of cytoplasmic dynein1 subunits or Adaptor Protein 2 (AP2) subunit. Our experiments confirmed the rapid degradation of these proteins and their functional inhibition in bulk cell populations. Additionally, to mitigate the reduced expression often associated with the degron tagging, we established a method to control expression levels by inserting a mini-promoter immediately upstream of the knock-in cassette. Our method simplifies the workflow for degron tag knock-ins and enhances the versatility of these valuable technologies.

## Introduction

Conditional protein depletion via degron-mediated degradation is a powerful tool for investigating protein function in complex biological systems and can also serve as a therapeutic approach to modulate disease-associated proteins ***Pillow et al. (2020); Maneiro et al. (2020)***. Conditional degron technologies have been developed based on proteolysis-targeting chimeras (PROTACs) ***Nabet et al. (2018, 2020); Buckley et al. (2015); Tovell et al. (2019)*** or molecular glue degraders ***Nishimura et al. (2009); Yesbolatova et al. (2020); Koduri et al. (2019); Sievers et al. (2018)*** that utilize E3 ubiquitin ligases, such as CRL4–Cereblon (CRBN) ***Nabet et al. (2018); Koduri et al. (2019); Sievers et al. (2018) and CRL2–Von Hippel-Lindau (VHL) Nabet et al. (2020); Buckley et al. (2015); Tovell et al. (2019)***. These include the degradation tag (dTAG) system ***Nabet et al. (2018)***, the Halo-PROTAC system ***Buckley et al. (2015)***, the Auxin-inducible degron (AID) ***Nishimura et al. (2009)***/AID2 ***Yesbolatova et al. (2020)*** system, and the IKAROS family zinc finger 3 degron (IKZF3d) system ***Sievers et al. (2018)***. These degrons can confer rapid degradation kinetics upon drug addition and reversible recovery of protein levels after the drug removal ***Nabet et al. (2018); Nishimura et al. (2009); Yesbolatova et al. (2020); Bondeson et al. (2022)***. To investigate the function of endogenous proteins, the degron tag technologies are often combined with the CRISPR-Cas9 system to knock in the degron tag at either the 5’ or 3’ end of the gene of interest ***Yesbolatova et al. (2020); Damhofer et al. (2021); Ghetti et al. (2021); Abuhashem and Hadjantonakis (2022); Kurashina and Mizumoto (2023)***. The CRISPR-Cas9 system induces precise double-strand breaks (DSB) in the targeted DNA via an RNA-mediated process ***Jinek et al. (2012)***, and the DSBs are repaired mainly by the two pathways: nonhomologous end-joining (NHEJ) and homology-directed repair (HDR) ***Scully et al. (2019)***. NHEJ often creates insertions or deletions at the repair site, whereas HDR utilizes a donor template that has homology to the sequence surrounding the DSB region for accurate repair and is employed for knocking in the degron tags. Although the HDR/NHEJ ratio is largely affected by the gene locus and cell types, NHEJ was shown to be predominant for repairing the Cas9-induced DSBs in human cells ***Miyaoka et al. (2016)***. The low HDR activity makes it difficult to obtain the homozygous degron tag knock-in cells, and laborious single-cell cloning is often considered necessary to select the cells that harbor degron tags in both alleles ***Damhofer et al. (2021); Abuhashem and Hadjantonakis (2022)***. This limitation may restrict applicable cell lines to those with relatively high HDR activity. Several strategies that can improve the HDR/NHEJ ratio have been developed. These include synchronizing the cell cycle phase to S and G2 ***Lin et al. (2014); Wienert et al. (2020); Yang et al. (2016)***, when HDR normally occurs ***Takata et al. (1998)***, inhibiting NHEJ ***Cloarec-Ung et al. (2024); Maruyama et al. (2015); Riesenberg and Maricic (2018)***, and enhancing HDR ***Charpentier et al. (2018); Pinder et al. (2015); Song et al. (2016); Zhang et al. (2020)***. However, those approaches may not yield consistent results ***Gerlach et al. (2018)*** or may lead to genomic instability ***Cullot et al. (2024)***.

Another consideration not frequently discussed is the alternation of the expression level of the protein of interest through the insertion of a degron tag. To efficiently inhibit the function, the expression level of the degron-tagged protein should be high enough to maintain its function before adding the drug, and low enough to compromise its function upon the addition of the drug. The previous systematic study demonstrated using the ectopic expression system that the expression level of the protein of interest is largely affected by the specific degron tag (FKBP12^F36V^, Halo-tag, AID/AID2, or IKZF3d) and the location (either N- or C-terminus) of the tag in the absence of the drug ***Bondeson et al. (2022)***. Similarly, the published protocol noted that the addition of FKBP12^F36V^ to the endogenous gene locus can lower the protein expression level in an unpredictable manner ***Damhofer et al. (2021)***. A versatile method to overcome this issue has not been established.

## Results

### The heterozygous dTAG knock-in system allows us to bypass single-cell cloning

We first determined the efficiency of obtaining homozygous degron tag knock-in in the immortalized retinal pigment epithelial cell (RPE1-hTERT), a genetically stable near-diploid cell ***Jiang et al. (1999)***, which is believed to have low HDR activity. The previous study required the utilization of two donor vectors that have different antibiotic resistance genes to obtain homozygous knock-in cells ***Miyamoto et al. (2015)***. Another study employed the NHEJ inhibitor to acquire a higher percentage of homozygous knock-in cells ***Ghetti et al. (2021)***. We transfected RPE1-hTERT cells expressing Blue Fluorescent Protein (BFP) tagged Cas9 ***Kanie et al. (2017)*** with a CRISPR RNA (crRNA) targeting near the start codon of a gene of interest (GOI) and a transactivating CRISPR RNA (tracrRNA) along with the donor cassette that consists of a left homology arm (LHA), a blasticidin resistance gene (BlastR), the self-cleaving peptides (P2A), V5-tag, FKBP12^F36V^, and a right homology arm (RHA) (**Figure 1A and Figure 1-figure supplement 1A-E**). The dTAG system utilizes FKBP12^F36V^ with the heterobifunctional degraders that bind to both FKBP12^F36V^ and the E3 ubiquitin ligases ***Nabet et al. (2018***, 2020). Upon addition of the degrader drug, the FKBP12^F36V^ fusion protein can be quickly degraded ***Nabet et al. (2018***, 2020). Following the blasticidin selection and single-cell sorting, we extracted genomic DNA from each single-cell clone and amplified the region flanking the crRNA targeting region using polymerase chain reaction (PCR) to test the insertion of the knock-in cassette. As expected, all the single-cell clones that we analyzed had knock-in alleles. The analyses of five knock-in lines revealed that almost all, except four, had the knock-in cassette in only one allele, confirming the low success rate of generating homozygous knock-in cells (**Figure 1B**).

**Figure 1.**
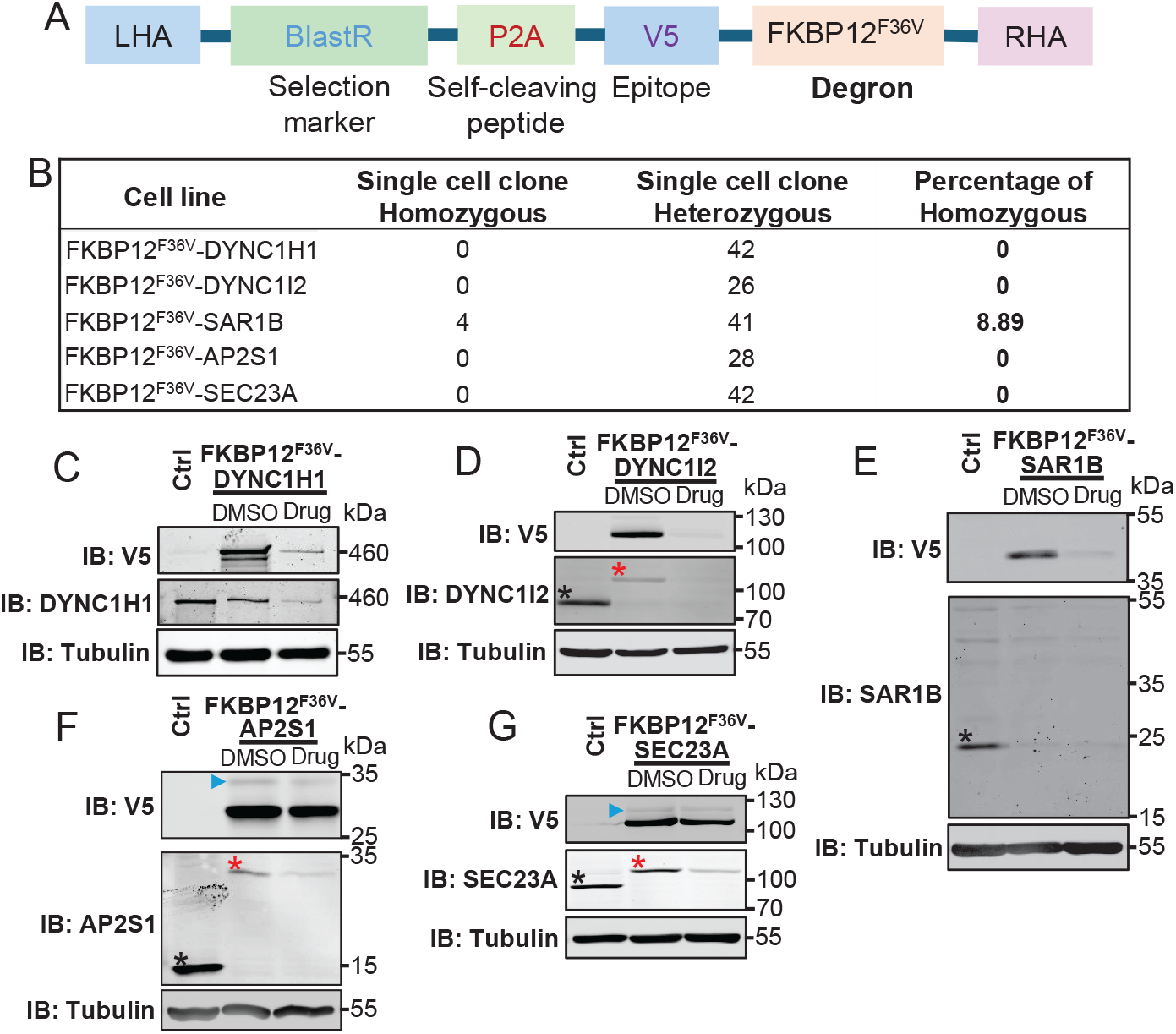
Generation and validation of the heterozygous knock-in cells harboring FKBP12^*F* 36*V*^ tagged proteins. **(A)** A diagram showing the design of the donor double-stranded DNA (dsDNA) for the FKBP12^*F* 36*V*^ knock-in cells. LHA, Left Homology Arm; BlastR, blasticidin resistance gene; RHA, Right Homology Arm. **(B)** A table showing the number of single-cell clone RPE-BFP-Cas9 knock-in cells that have homozygous or heterozygous knock-in alleles and the percentage of homozygous knock-in cells. PCR amplified DNA surrounding the guide RNA targeting region of each Gene of Interest (GOI) was analyzed through DNA electrophoresis. **(C–G)** Immunoblot (IB) analysis of the indicated proteins in the heterozygous knock-in RPE-BFP-Cas9 cells expressing GOI fused to V5 and FKBP12^*F* 36*V*^. Cells grown to confluence were treated with either dimethyl sulfoxide (DMSO) or 1 μM dTAG-13 (drug) for 24 hours. Tubulin (IB: Tubulin) serves as a loading control. Molecular weights (kDa) estimated from a protein marker are indicated. The black asterisks indicate the endogenous proteins; The red asterisks indicate the fusion proteins; The blue arrowheads indicate the slower-migrating fusion proteins, which likely come from the ribosome read-through of the P2A peptide (i.e., blasticidin resistance gene-V5-FKBP12^*F* 36*V*^ fusion proteins).

To overcome the limitation of efficiently obtaining homozygous degron tag knock-in cells, we sought to develop a method that allows us to use heterozygous cells rather than boosting the chance of obtaining homozygous cells by increasing the HDR/NHEJ ratio. We predicted that when the degron cassette is introduced by HDR following a double-strand break induced by Cas9, the other allele may also be excised by the enzyme and repaired through NHEJ, leading to an insertion/deletion in that allele. To test our hypothesis, we analyzed the genomic DNA sequence of our bulk knock-in cells before the single-cell sorting using the Tracking of Indels by DEcomposition (TIDE) analysis ***Brinkman et al. (2014)***, which allows us to analyze the frequency of insertions/deletions in a mixed cell population. The analysis confirmed that the allele that did not have the knock-in cassette had either frame-shift insertion/deletion or its start codon was deleted in almost all cells in four out of five knock-in lines that we tested (**Figure 1-figure supplement 2A-E**).

As an exception, FKBP12^F36V^-SAR1B knock-in lines showed 96% of the non-frame shifting minus 3 deletions (**Figure 1-figure supplement 2C**). To validate our knock-in bulk cells at the protein level and assess degradation efficiency across different Proteins of Interest (POIs), we performed immunoblots of the bulk knock-in cells using antibodies against the V5 epitope tag or the endogenous proteins (**Figure 1C-G**). Consistent with the genomic DNA analyses, the bands that correspond to the endogenous proteins were almost undetectable in the bulk knock-in cell lines (black asterisks in **Figure 1D-G**), suggesting that most of the heterozygous knock-in cells express only the degron fusion proteins. While the genomic DNA analyses showed the dominant 3bp in-frame deletion (**Figure 1-figure supplement 2C**), the expression of the endogenous SAR1B protein was greatly diminished in the FKBP12^F36V^-SAR1B knock-in line (**Figure 1E**). Upon 24-hour treatment with the degrader drug, dTAG-13 ***Nabet et al. (2018)***, each FKBP12^F36V^ fusion protein was potently degraded to varying extents (**Figure 1C-G**). These results indicate that the dTAG system is effective in our bulk heterozygous knock-in cell lines, thereby enabling the bypassing of laborious single-cell cloning.

### Comparison of three different degron tag systems in our heterozygous knock-in system

To verify the applicability of our new heterozygous knock-in system to different degron tag systems and compare the efficiency of the different systems, we expanded our focus to three popular degrons: FKBP12^F36V^ ***Nabet et al. (2018)***, IKZF3d (IKZF3 aa130–189) ***Sievers et al. (2018)***, and HaloTag ***Buckley et al. (2015)***. The three degrons utilize different degradation mechanisms (**Figure 1-figure supplement 2F**). The compounds dTAG-13 and pomalidomide (an IKZF3d degrader) recruit CRBN ***Nabet et al. (2018); Sievers et al. (2018)***, whereas the HaloPROTAC3 (a HaloTag degrader) binds to VHL ***Buckley et al. (2015)***. We generated a panel of nine knock-in bulk cell lines for three degrons, each crossed with three POIs, SAR1B, AP2S1, and SEC23A (**Figure 2A**). Immunoblot analyses showed that the endogenous proteins were efficiently removed in all three systems (**Figure 2B-D**), except for FKBP12^F36V^-SAR1B (**Figure 2B**), which harbors a non-frameshifting 3 bp deletion (**Figure 1-figure supplement 2C**). While a band slightly upper-shifted than the endogenous AP2S1 was detected in the knock-in cells that harbor HaloTag-AP2S1 (blue asterisk in **Figure 2C**), genomic DNA analyses showed that the allele mainly has “frame-shifting mutations” (**Figure 2-figure supplement 1A**). Consistent with the previous study ***Bondeson et al. (2022)***, which used an overexpression system to compare different degron tags, our analyses showed that the basal expression level of the degron tag fusion proteins and their degradation efficiency upon the drug treatment greatly vary among the different degrons (**Figure 2B-D**). Interestingly, we consistently observed the expression level to be the highest with the HaloTag fusions, the lowest with the IKZF3d fusions, and the median with the FKBP12^F36V^ fusions. Notably, the expression levels of the fusion proteins are generally lower than those of endogenous proteins (compare endogenous proteins marked by black asterisks with the fusion proteins marked by red asterisks in **Figure 2C and D**), and SAR1B fusion proteins are undetectable with an antibody against the protein (**Figure 2B**). The degradation efficiency tended to be the highest with the IKZF3d fusions, and the HaloTag fusions had a slightly higher degradation rate than the FKBP12^F36V^ fusions (**Figure 3-figure supplement 1 and 2**). Additionally, we occasionally observed upward-shifted bands (blue arrowheads in **Figure 1F-G and Figure 2C-D**), which likely correspond to blasticidin-resistant gene fusion proteins resulting from ribosome read-through of the P2A peptide ***Liu et al. (2017)***. The degradation efficiency of the upshifted bands varied among the fusion proteins.

**Figure 2.**
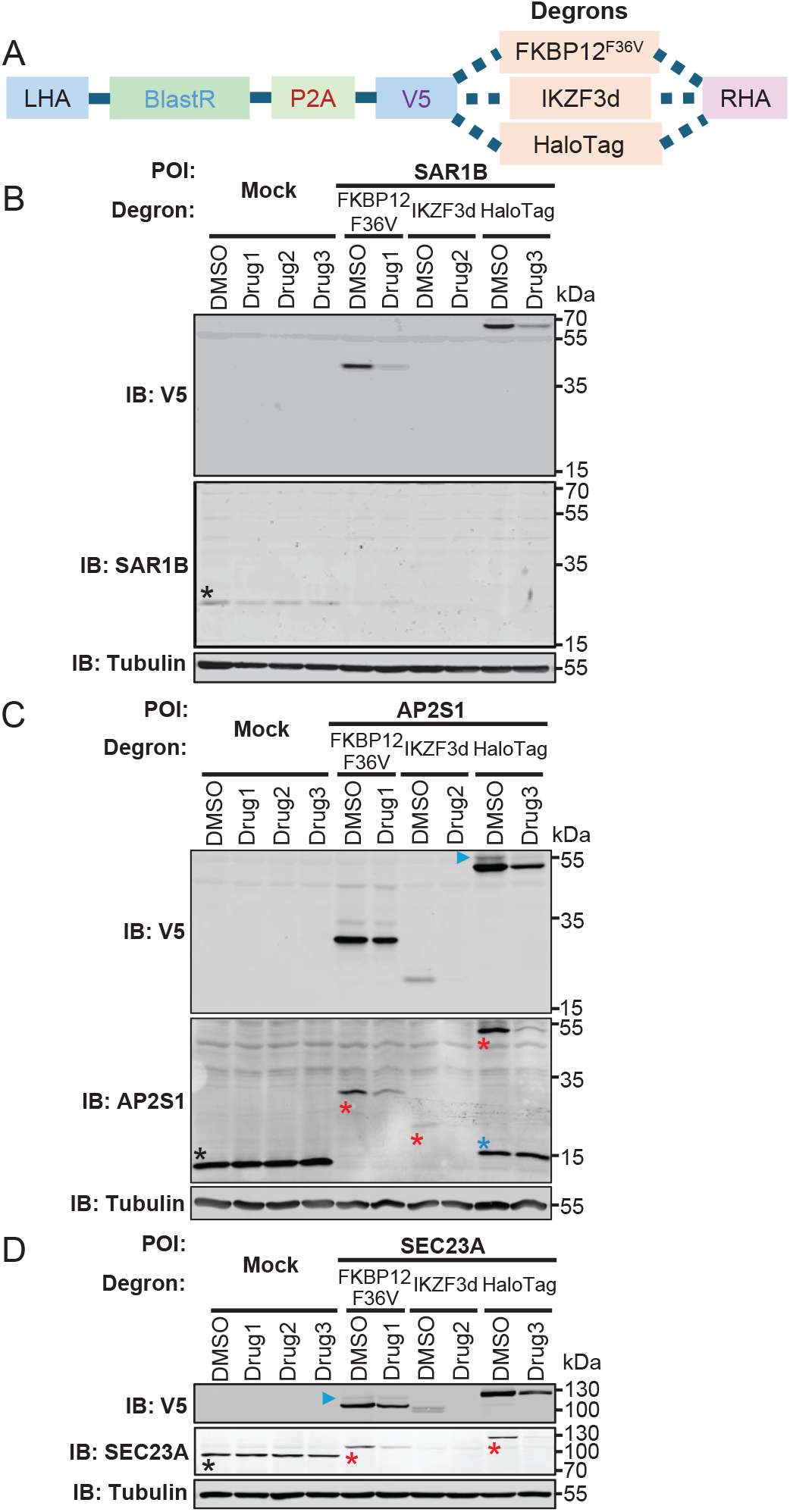
Generation and validation of the heterozygous knock-in cells harboring different degron-tagged proteins. **(A)** A diagram illustrating the design of the donor double-stranded DNA (dsDNA) used for the generation of knock-in cells that have three different degron tags. The three degrons are FKBP12^*F* 36*V*^ (for dTAG system), IKZF3d (IKZF3 aa130–189), and HaloTag7. **(B-D)** Immunoblot (IB) analysis of the indicated proteins in the heterozygous knock-in RPE-BFP-Cas9 cells expressing GOI fused to V5 and indicated degron (FKBP12^*F* 36*V*^, IKZF3d, and HaloTag). The cells grown to confluence were treated with either dimethyl sulfoxide (DMSO) or 1 μM of the indicated drug (drug1: dTAG-13, drug2: pomalidomide, and drug3: HaloPROTAC3) for 24 hours. Tubulin (IB: Tubulin) serves as a loading control. Molecular weights (kDa) estimated from a protein marker are indicated. The black asterisks indicate the endogenous proteins; The red asterisks indicate the fusion proteins; The blue asterisk indicates the band slightly shifted to the upper region than the endogenous AP2S1. The blue arrowheads indicate the slower-migrating fusion proteins, which likely come from the ribosome read-through of the P2A peptide (i.e., blasticidin resistance gene-V5-FKBP12^*F* 36*V*^ fusion proteins).

**Figure 3.**
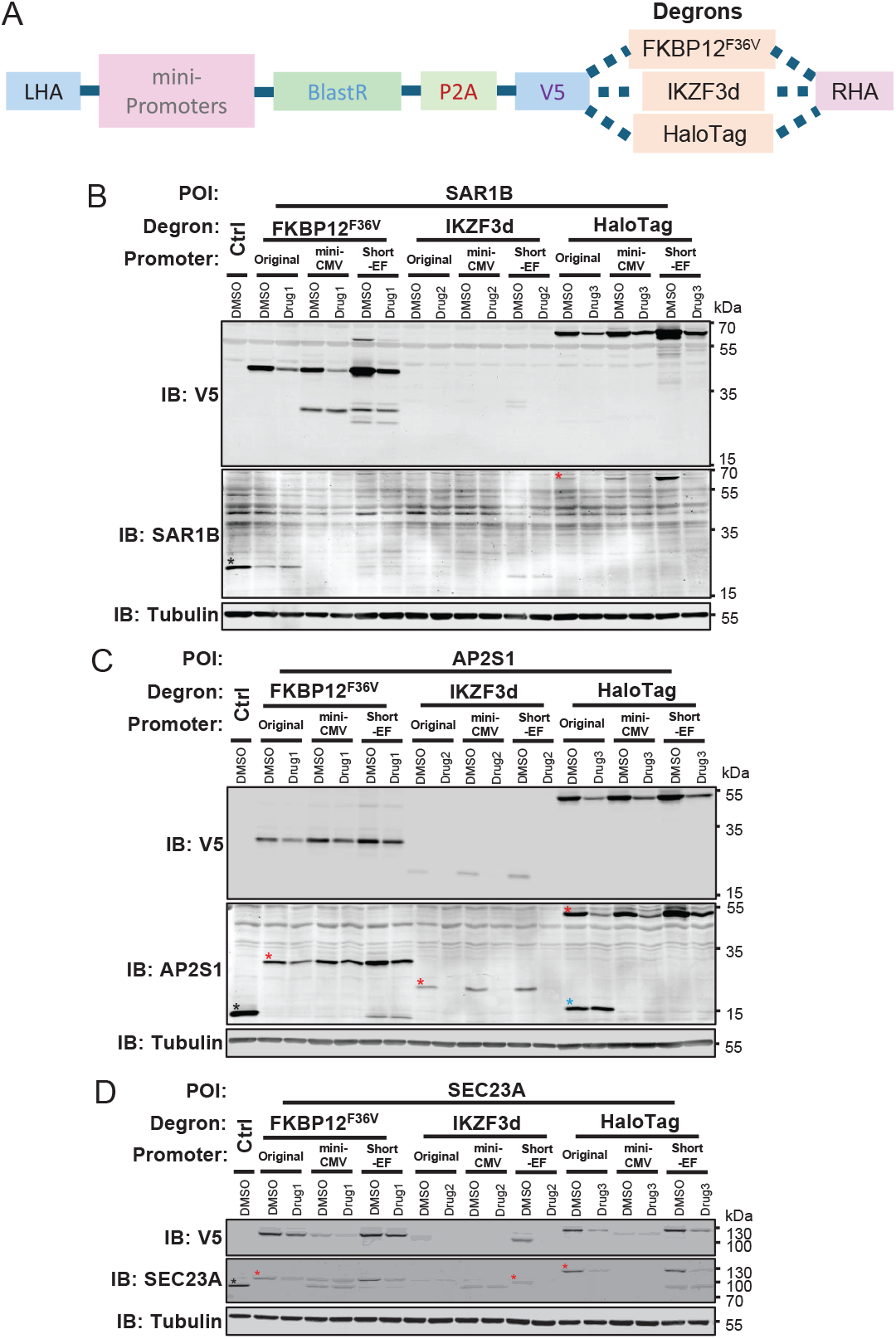
Generation and validation of the heterozygous knock-in cells that express degron-tagged proteins regulated by different mini-promoters. **(A)** A diagram showing the design of the donor double-stranded DNA (dsDNA) to incorporate the mini-promoter (mini-CMV promoter and short-EF promoter) in the degron-tag knock-in cells. **(B-D)** Immunoblot (IB) analysis of the indicated proteins in the heterozygous knock-in RPE-BFP-Cas9 cells that express the gene of interest fused to V5 and indicated degron (FKBP12^*F* 36*V*^, IKZF3d, and HaloTag under the control of the original promoter, mini-CMV promoter, or the short-EF promoter. The cells grown to confluence were treated with either dimethyl sulfoxide (DMSO) or 1 μM indicated drug (drug1: dTAG13, drug2: pomalidomide, and drug3: HaloPROTAC3) for 24 hours. Tubulin (IB: Tubulin) serves as a loading control. Molecular weights (kDa) estimated from a protein marker are indicated. The black asterisks indicate the endogenous proteins; The red asterisks indicate the fusion proteins; The blue asterisk indicates the band slightly shifted to the upper region than the endogenous AP2S1. Quantification data are available in **Figure 3-figure supplement 1 and 2**.

### Modulation of the expression level of the degron fusion proteins through mini promoters

As described above, we often observed lower expression levels of the degron fusion proteins than those of endogenous proteins (**Figure 2B-D**). The lower expression may compromise the function of the proteins even before adding the degrader drugs. Therefore, we sought to develop an approach to modulate the expression of the fusion proteins, so that they can retain the function until the degradation is induced through the degrader drugs. To do this, we incorporated either an enhancer-less core sequence (from −71 to −1) of the cytomegalovirus promoter ***Boshart et al. (1985)*** (mini-CMV) or a promoter derived from the human EF-1 gene that lacks the first intron ***Kim et al. (1990)*** (short-EF) in front of the BlastR gene in the dsDNA donor cassette (**Figure 3A**). Including the original constructs without the exogenous promoters (**Figure 2A**), we generated a panel of nine heterozygous knock-in cell lines (three promoters cross three degrons) for each of the three different POIs. Immunoblot analyses showed that endogenous proteins were eliminated in 21 out of 27 cell lines (**Figure 3B-D**). Endogenous proteins were still expressed in the remaining six cell lines, likely because most of these cells have in-frame insertion/deletion (**Figure 1-figure supplement 2C and Figure 3-figure supplement 3**). The assay also revealed that the integration of short-EF promoter consistently provided higher expression than the constructs without the exogenous promoters, whereas enhancement of expression by the mini-CMV promoter was less pronounced and depended on the tags and POIs (compare “original” vs “mini-CMV” in each tag and POI in **Figure 3B-D, Figure 3-figure supplement 1A-C, and Figure 3-figure supplement 2A-C**). Notably, with the addition of a short-EF promoter, we were able to bring the expression level of SAR1B fused with HaloTag to a level closer to the endogenous one (**Figure 3B and Figure 3-figure supplement 2A**). Even with the short-EF promoter, we failed to achieve a comparable expression level to the endogenous proteins with the IKZF3d fusion proteins (**Figure 3B-D and Figure 3-figure supplement 2A-C**). When we added the degrader drugs, the expression levels of the fusion proteins regulated by the exogenous promoters diminished to a similar extent as those without the mini-promoters in all the fusion proteins that we tested (**Figure 3B-D, Figure 3-figure supplement 1A-C, and Figure 3-figure supplement 2A-C**).

Together, the data suggest that we can control the basal expression levels of the degron fusion proteins through the mini-promoters without affecting their drug-induced degradation efficiency.

### Functional validation of the heterozygous degron knock-in cells

Conditional protein degradation allows for precise and controlled analysis of essential genes, enabling researchers to understand cellular functions by rapidly inducing the degradation of specific proteins. The kinetics of degradation can vary and may be unpredictable, depending on both the degron and the target proteins involved ***Bondeson et al. (2022)***. To test if the degrader drugs induce rapid removal of the proteins, and thus inhibit the function of the target proteins in our heterozygous knock-in cells, we chose three proteins that are known to be essential for cell division or growth: DYNC1H1 ***Raaijmakers et al. (2013)***, DYNC1I2 ***Raaijmakers et al. (2013)***, and AP2S1 ***Zimmerman et al. (2025)*** (**Table 1**).

**Table 1.** The abbreviation of the POIs used in this study.

| GOI | Name | MW (kDa) | UniprotID |
| --- | --- | --- | --- |
| DYNC1H1 | dynein cytoplasmic 1 heavy chain 1 | 532 | Q14204 |
| DYNC1I2 | dynein cytoplasmic 1 intermediate chain 2 | 68 | Q8TCX1 |
| SAR1B | secretion-associated ras-related GTPase 1B | 22 | Q9Y6B6 |
| SEC23A | SEC23 homolog A, COPII component | 82 | Q15436 |
| AP2S1 | adaptor related protein complex 2 subunit sigma 1 | 17 | P53680 |

Cytoplasmic dynein 1, a minus-end-directed motor complex, is essential for mitosis. It plays crucial roles in mitotic spindle organization ***Vaisberg et al. (1993); Faulkner et al. (2000)***, chromosome movement ***Sharp et al. (2000)***, and kinetochore protein transport ***Howell et al. (2001)***. It is also important for maintaining the structure and the localization of the Golgi apparatus ***Harada et al. (1998); Palmer et al. (2009)***. Depletion of dynein subunits, such as DYNC1H1 or DYNC1I2, has been shown to cause mitotic arrest ***Raaijmakers et al. (2013)*** and the dispersal of the Golgi apparatus ***Harada et al. (1998); Palmer et al. (2009)***.

We used the cells harboring either DYNC1H1 or DYNC1I2 fused with FKBP12^F36V^ (**Figure 1C and D**) as model cases to evaluate degradation kinetics and functional defects upon addition of the degrader drug, dTAG-13, in our heterozygous knock-in cell lines. We found that DYNC1H1 and DYNC1I2 degradation began within 1 hour after administering the degrader drug and reached the maximum efficiency within 3 hours for the DYNC1H1-fusions (**Figure 4A and B**) and 6 hours for the DYNC1I2-fusions (**Figure 5A and B**), respectively. The degradation effect persisted for 24 hours for both fusion proteins without replacing the media (**Figure 4A-B and 5A-B**). Next, we investigated the impact of dynein1 subunit depletion on mitotic progression and Golgi positioning. Although the expression level of the fusion proteins was 50% or 35% for DYNC1H1 and DYNC1I2 before adding the degrader drugs, consistent with the cells having only one knock-in allele (heterozygous), the cells did not show apparent mitotic arrest or the Golgi dispersal (**Figure 4-figure supplement 1A and B**). This suggests that the cells can still retain the function of cytoplasmic dynein with the reduced expression of the subunits and that the fusion proteins are functionally intact. Upon addition of the drug, we started to see a statistically significant increase in the mitotic index at 6 hours, and the percentage of the cells arrested at mitosis increased over the time course of drug treatment in both knock-in cell lines (**Figure 4C-D** and **Figure 5C-D**). Concurrently, the total number of cells per micrograph decreased over time in the cells treated with the degrader drug, suggesting that the cells depleted of DYNC1H1 likely detach from the plate following the mitotic arrest (**Figure 4E**). The total number of cells did not significantly decrease in the cells depleted of DYNC1I2 (**Figure 5E**), likely reflecting the milder effect of DYNC1I2 depletion compared with the DYNC1H1 removal. When we analyzed the Golgi structure using an anti-TGN46 antibody, which recognizes the Trans-Golgi network ***Lujan et al. (2024)***, we observed Golgi dispersal within 6 hours of the drug treatment in the DYNC1H1 line (**Figure 4G**); Golgi dispersal was observed with 3 hours of drug treatment in the DYNC1I2 line (**Figure 5G**). This suggests a more pronounced effect of DYNC1I2 depletion compared with the DYNC1H1 removal, likely due to the direct interaction of the dynein1 intermediate chain with the golgin160 for the Golgi positioning ***Yadav et al. (2012)***. These data suggest that the function of the dynein subunits is intact before the drug treatment and is quickly inhibited following the drug administration in both heterozygous knock-in cells. In contrast, the degrader drug dTAG-13 did not visibly affect the mitotic cell count or the Golgi structure of the parental cells (**Figure 4-figure supplement 1A-B**).

**Figure 4.**
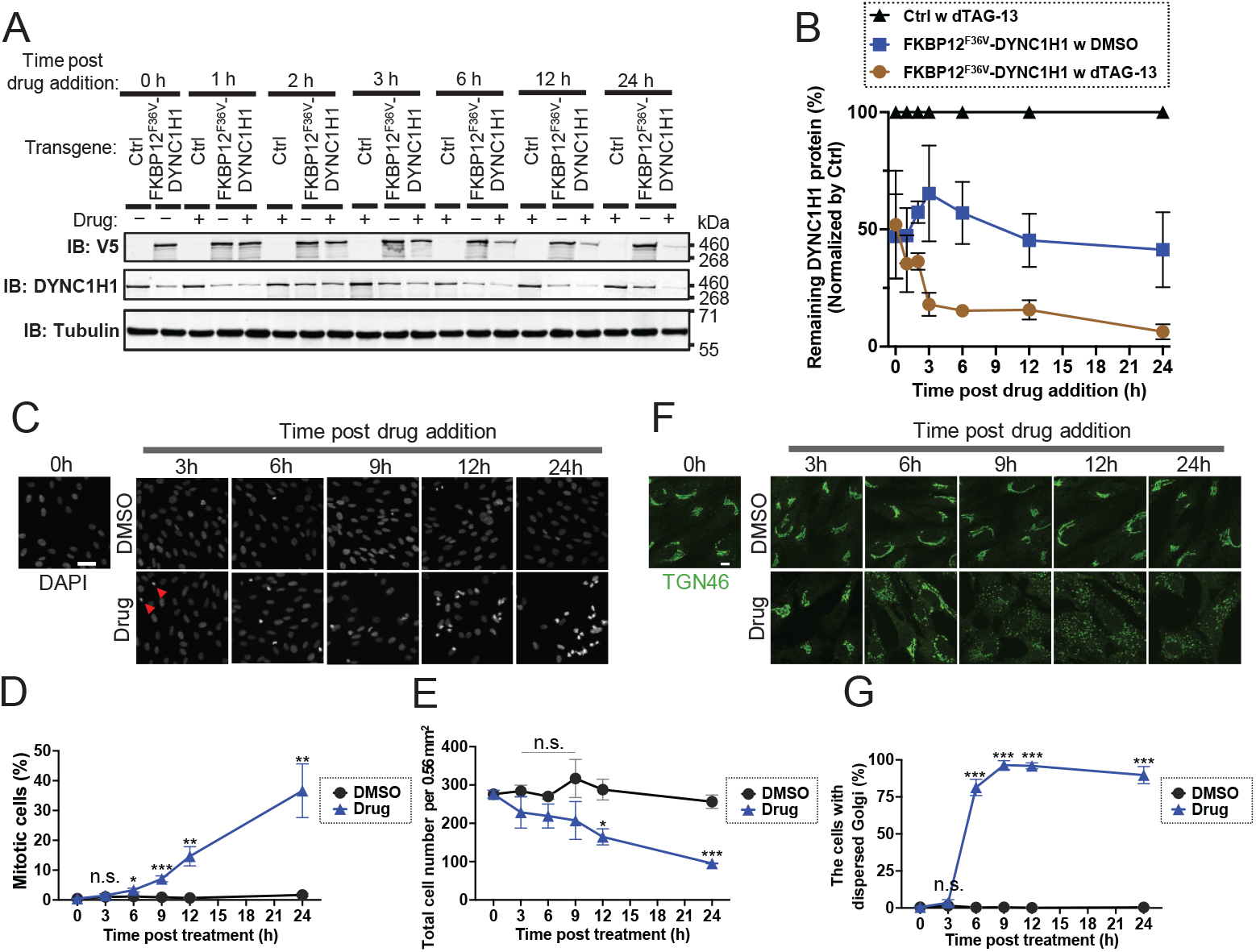
The degradation kinetics and function assay of cytoplasmic dynein1 in the heterozygous knock-in cells expressing FKBP12^*F* 36*V*^-DYNC1H1. **(A)** Immunoblot (IB) analysis of V5 (IB: V5) and DYNC1H1 (IB: DYNC1H1) in the heterozygous knock-in RPE-BFP-Cas9 cells expressing FKBP12^*F* 36*V*^ tagged DYNC1H1. The cells grown to confluence were treated with either dimethyl sulfoxide (DMSO) or 1 μM dTAG-13 (drug) for the indicated time. Tubulin (IB: Tubulin) serves as a loading control. Molecular weights (kDa) estimated from a protein marker are indicated. **(B)** Quantification of the DYNC1H1 protein level shown in **Figure 4A**. DYNC1H1 protein level (%) was calculated by normalizing the expression level at each time point to that of the control cells at the same time point. The data were averaged from three independent experiments. Error bars represent Mean ± SEM. **(C)** Mitosis arrest assay in the heterozygous knock-in RPE-BFP-Cas9 cells expressing FKBP12^*F* 36*V*^ tagged DYNC1H1. The cells were treated with either dimethyl sulfoxide (DMSO) or 1 μM dTAG-13 (drug) for the indicated time, fixed, and stained with DAPI. The representative image from each condition is shown. The red arrowheads indicate the mitotic cells. Scale bar: 50 μm **(D)** Quantification of the percentage of the mitotic cells shown in **Figure 4C**. The data were averaged from three independent experiments. Error bars represent ± SEM. Statistics obtained through comparing the drug-treated cells and the control at each time point by multiple unpaired t-tests. **(E)** Quantification of the total cell number of the cells per microscope image (0.56 mm^2^) from the experiments shown in **Figure 4C**. The data were averaged from three independent experiments. Error bars represent ± SEM. Statistics obtained through comparing the drug-treated cells and the control at each time point by multiple unpaired t-tests. **(F)** The Golgi dispersal assay in the heterozygous knock-in RPE-BFP-Cas9 cells expressing FKBP12^*F* 36*V*^ tagged DYNC1H1. The cells were treated with either dimethyl sulfoxide (DMSO) or 1 μM dTAG-13 (drug) for the indicated time, fixed, and stained with anti-TGN46 antibody. Scale bar: 10 μm **(G)** Quantification of the percentage of the cells with dispersed Golgi in the cells shown in **Figure 4F**. The data were averaged from three independent experiments. Error bars represent ± SEM. Statistics obtained through comparing the drug-treated cells and the control at each time point by multiple unpaired t-tests. n.s., not significant; *p < 0.05, **p < 0.01, ***p < 0.001

**Figure 5.**
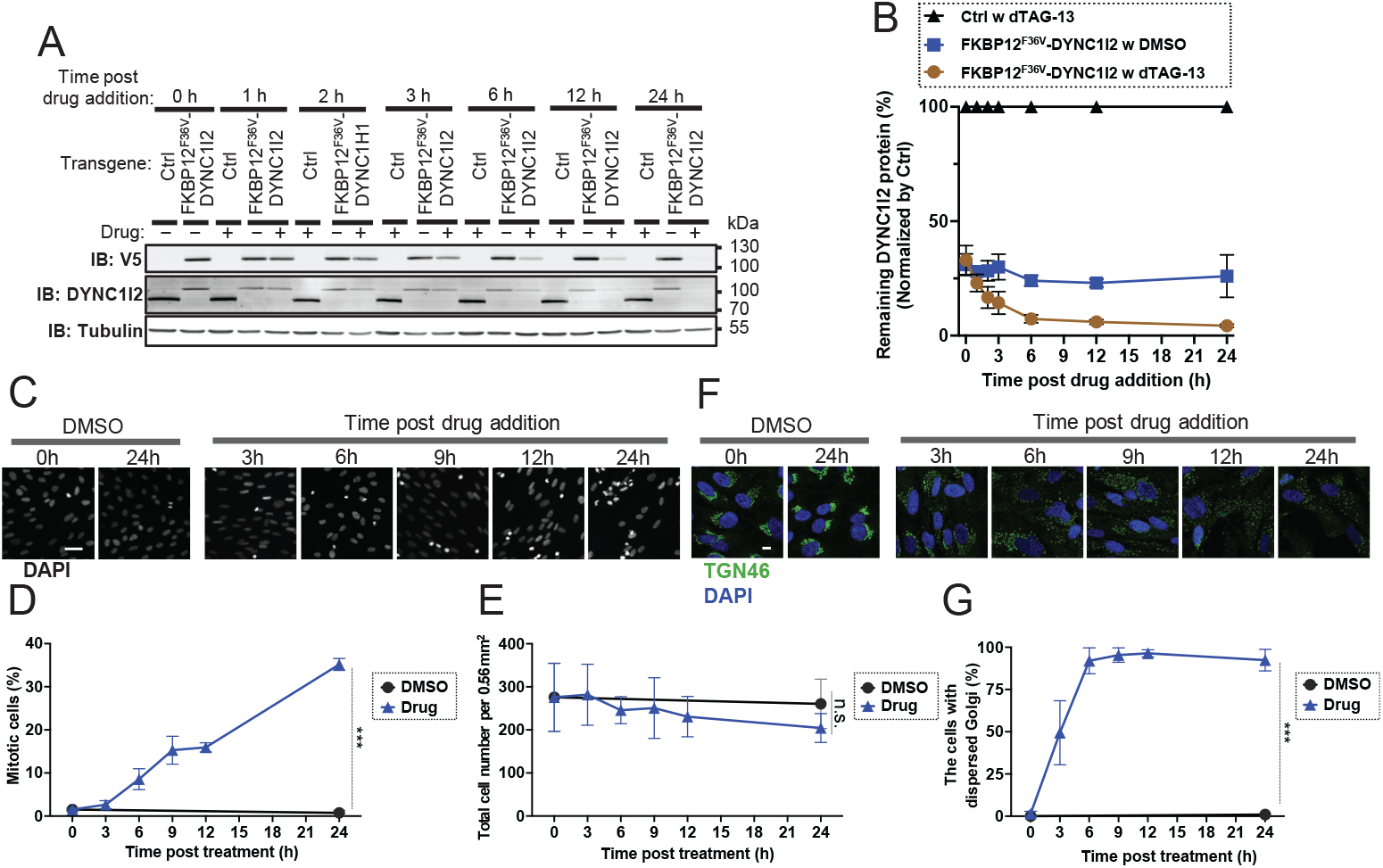
The degradation kinetics and function assay of cytoplasmic dynein1 in the heterozygous knock-in cells expressing FKBP12^*F* 36*V*^-DYNC1I2. **(A)** Immunoblot (IB) analysis of V5 (IB: V5) and DYNC1I2 (IB: DYNC1I2) in the heterozygous knock-in RPE-BFP-Cas9 cells expressing FKBP12^*F* 36*V*^ tagged DYNC1I2. The cells grown to confluence were treated with either dimethyl sulfoxide (DMSO) or 1 μM dTAG-13 (drug) for the indicated time. Tubulin (IB: Tubulin) serves as a loading control. Molecular weights (kDa) estimated from a protein marker are indicated. **(B)** Quantification of the DYNC1I2 protein level shown in **Figure 5A**. DYNC1I2 protein level (%) was calculated by normalizing the expression level at each time point to that of the control cells at the same time point. The data were averaged from three independent experiments. Error bars represent Mean ± SEM. **(C)** Mitosis arrest assay in the heterozygous knock-in RPE-BFP-Cas9 cells expressing FKBP12^*F* 36*V*^ tagged DYNC1I2. The cells were treated with either dimethyl sulfoxide (DMSO) or 1 μM dTAG-13 (drug) for the indicated time, then fixed and stained with DAPI. The representative image from each condition is shown. Scale bar: 50 μm **(D)** Quantification of the percentage of the mitotic cells shown in **Figure 5C**. The data were averaged from three independent experiments. Error bars represent ± SEM. Statistics obtained through comparing the drug-treated cells and the control at 24 hours by an unpaired t-test. **(E)** Quantification of the total cell number of the cells per microscope image (0.56 mm^2^) from the experiments shown in **Figure 5C**. The data were averaged from three independent experiments. Error bars represent ± SEM. Statistics obtained through comparing the drug-treated cells and the control at 24 hours by an unpaired t-test. **(F)** The Golgi dispersal assay in the heterozygous knock-in RPE-BFP-Cas9 cells expressing FKBP12^*F* 36*V*^ tagged DYNC1I2. The cells were treated with either dimethyl sulfoxide (DMSO) or 1 μM dTAG-13 (drug) for the indicated time, fixed, and stained with anti-TGN46 antibody and DAPI. Scale bar: 10 μm **(G)** Quantification of the percentage of the cells with dispersed Golgi in the cells shown in **Figure 5F**. The data were averaged from three independent experiments. Error bars represent ± SEM. Statistics obtained through comparing the drug-treated cells and the control at 24 hours by an unpaired t-test. n.s., not significant; *p < 0.05, **p < 0.01, ***p < 0.001

As a second example, we tested the ability of the heterozygous knock-in cells that harbor HaloTag-AP2S1 to recapitulate the previously reported effect of AP2S1 loss on endocytosis ***Xu et al. (2024)***. AP2S1 is a subunit of the adaptor protein complex 2 (AP-2) ***Collins et al. (2002)***, a critical regulator for clathrin-mediated endocytosis ***Motley et al. (2003)***. AP2 binds to membrane protein cargos, initiates the assembly of clathrin-coated pits, and promotes internalization of the cargo proteins, which include transferrin receptors ***Xu et al. (2024); Motley et al. (2003)***. The loss of AP2S1 was shown to destabilize other subunits ***Zimmerman et al. (2025); Li et al. (2025)*** and inhibit endocytosis of the transferrin receptors ***Xu et al. (2024)***. In the heterozygous knock-in cell lines expressing HaloTag-AP2S1 under the control of the original promoter, the degradation of AP2S1-fusions induced by HaloPROTAC3, a degrader drug for HaloTag proteins, began one hour after degrader administration, reached its 90% degradation efficiency at three hours, and the effects of degradation lasted for 24 hours (**Figure 6A and B**). We then performed the transferrin internalization assay at 6 hours following the drug administration using the spinning disk confocal microscopy. In the cells treated with DMSO, the dotty patterns of fluorescent transferrin were readily observed in the cytoplasm (**Figure 6C and D**), indicating that transferrin was successfully internalized via endocytosis in the assay condition. In contrast, we could not detect similar dotty staining in the cytoplasm of the majority of the drug-treated cells (**Figure 6C and D**), suggesting that endocytosis was efficiently inhibited by AP2S1 depletion. To better visualize internalization of transferrin, we conducted the optical reassignment-based super-resolution imaging ***Azuma and Kei (2015)*** to provide a 3D view of the fluorescent transferrin signals. The axial view of the 3D reconstructed images more clearly showed that most of the fluorescent-transferrin signals were detected in the cytoplasm in DMSO-treated cells, whereas the signals remained largely on the plasma membrane in the drug-treated cells (**see xz and yz views in the Figure 6E**). Notably, when we analyzed another cell line that harbors AP2S1 fusion proteins, short-EF promoter-regulated FKBP12^F36V^-AP2S1, the effect of the degrader drugs on endocytosis of transferrin was less pronounced (**Figure 6-figure supplement 1A-B**). The weaker phenotypes of those cells likely reflect the milder degradation efficiency of the FKBP12^F36V^-AP2S1 fusion protein than the HaloTag-AP2S1 fusion (**Figure 3C and Figure 3-figure supplement 2B**). This highlights the importance of carefully evaluating both expression level and functional effects upon the degrader drug treatment for each POI fusion.

**Figure 6.**
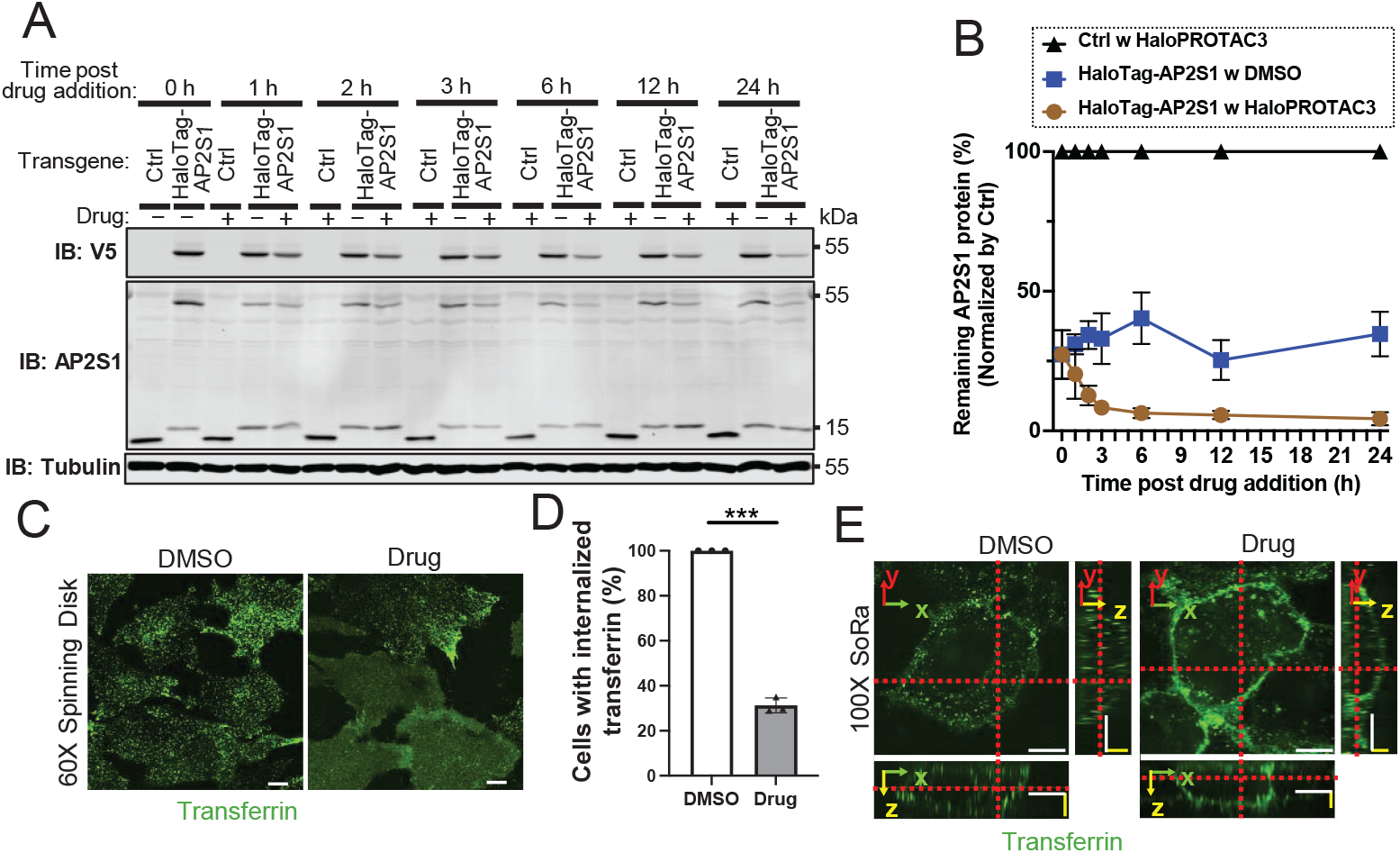
The degradation kinetics and function assay of AP2S1 in the heterozygous knock-in cells expressing HaloTag-AP2S1. **(A)** Immunoblot (IB) analysis of V5 (IB: V5) and AP2S1 (IB: AP2S1) in the heterozygous knock-in RPE-BFP-Cas9 cells expressing HaloTag-AP2S1. The cells grown to confluence were treated with either dimethyl sulfoxide (DMSO) or 1 μM HaloPROTAC3 (drug) for the indicated time. Tubulin (IB: Tubulin) serves as a loading control. Molecular weights (kDa) estimated from a protein marker are indicated. **(B)** Quantification of the AP2S1 protein level shown in **Figure 6A**. AP2S1 protein level (%) was calculated by normalizing the expression level at each time point to that of the control cells at the same time point. The data were averaged from three independent experiments. Error bars represent Mean ± SEM. **(C)** Transferrin internalization assays in the heterozygous knock-in RPE-BFP-Cas9 cells expressing HaloTag-AP2S1. The cells were treated with either dimethyl sulfoxide (DMSO) or 1 μM HaloPROTAC3 (drug) for 6 hours and incubated with transferrin-Alexa Fluor 488. Live-cell imaging was performed on a spinning disk confocal microscope. The individual image is from a representative z-slice. Scale bar: 10 μm **(D)** Quantification of the percentage of the cells with internalized transferrin shown in **Figure 6C**. The data were averaged from three independent experiments. Error bars represent ± SEM. Statistics obtained through comparing the drug-treated cells and the control by an unpaired t-test. ***p < 0.001 **(E)** 3D Super-resolution live-cell imaging was used to assess transferrin internalization assays in the condition described in **Figure 6C**. The representative 100x SoRa super-resolution z-slice images with orthogonal views are shown. For easier visualization, we have intensively enlarged the z-thickness of the orthogonal views. Scale bar for the x & y-axis: 10 μm; scale bar for the z-axis: 2 μm.

Nevertheless, all the functional validation tests confirmed that the heterozygous knock-in cells generated through our strategy can be used to test the function of essential genes.

## Discussion

Conditional degron tag systems are important tools to investigate the function of proteins through rapid degradation of them; however, two major issues restrict their application: 1) difficulty in obtaining homozygous knock-in cells that express degron tag fusion proteins, and 2) adding a degron tag may alter the expression level of the protein even in the absence of the degrader drug. In the current work, we confirm the presence of the two problems through the generation of a good number of the degron-tag knock-in cells. Only 4 of 183 single-cell clones that we analyzed had the knock-in cassette in both alleles (**Figure 1B**), indicating the low yield of homozygous knock-in cells. While the extent varied across different degron tags and proteins of interest, the expression levels of the degron-tag fusion proteins were often lower than those of endogenous proteins (**Figure 1C-G and 2B-D**). Reduced expression of fusion proteins has been reported even in knock-in cells that express the fusion proteins from their endogenous gene locus ***Cho et al. (2022); Taylor et al. (2025)***, likely because foreign tags alter mRNA stability, protein folding, and protein interaction. To mitigate these problems and improve the applicability of these valuable tools, we report here a strategy to efficiently and rapidly obtain the degron tag knock-in cells that can be used for functional assays. Our method was established based on the observation that the allele that does not have the knock-in cassette is edited by Cas9 in virtually all the cells that we analyzed, and the majority of the non-knock-in alleles had frame-shift insertion/deletion (**Figure 1-figure supplement 2 and Figure 2-figure supplement 1**). Consistent with the frame-shift insertion/deletion, immunoblot analyses failed to detect the bands that correspond to the endogenous proteins in 23 out of 29 knock-in cells that we created (**Figure 1C-D and Figure 3A-C**). This allowed us to skip single-cell cloning and use the bulk cells for functional assays. Subsequent analyses confirmed that the function of the target proteins is rapidly and efficiently inhibited upon addition of the degrader drugs in bulk knock-in cells (**Figures 4-6**). We also introduce a method to mitigate the reduction in the basal expression level of degron-tagged POIs by inserting minimal promoters at the beginning of the donor cassette (**Figure 3**). We tested two different mini-promoters and found that the short-EF promoter consistently provides higher expression than the one without mini-promoters, whereas the effect of minimal CMV varies among POIs and degron tags **(Figure 3, Figure 3-figure supplement 1 and 2)**.

While our approach has multiple advantages over commonly used methods that select homozygous clones for degron tag knock-in, there are several potential issues. In the subsequent sections, we will discuss the pros and cons of our methodology and a possible strategy that can be used for further improvement.

### Efficiency of obtaining homozygous knock-in cells

Consistent with the previously reported difficulty in obtaining RPE cells that harbor a knock-in cassette in both alleles ***Ghetti et al. (2021); Miyamoto et al. (2015)***, our results showed that the chances of gaining homozygous knock-in cells are very low (2.2% as shown in **Figure 1B**). In contrast, a recent study reported a much higher percentage of homozygous knock-in RPE cells, while the efficiency is affected by GOIs and crRNA sequences ***Philip et al. (2025)***. There are several methodological differences in the generation of knock-in cells between our report and the previous one. To create knock-in cells, we transfected RPE cells stably expressing BFP-Cas9 with a 1:500 ratio of donor dsDNA/guide RNA, whereas the previous study co-transfected RPE cells with a vector containing both Cas9 and guide RNAs with a plasmid carrying a donor template. Several studies suggest that the stoichiometry ratio of Cas9, guide RNA, and donor dsDNA is an essential factor influencing HDR-mediated knock-in efficiency ***Lin et al. (2014); Shirk et al. (2026); Chenouard et al. (2023); Schumann et al. (2015)***. While the intracellular concentrations of those components are difficult to predict in our current approach, stoichiometry optimization may further enhance HDR activity and improve the efficiency of successfully generating knock-in cells. Another notable difference is the size of the 5’ and 3’ homology arms. We adopted the previously established protocol, which uses 40 base pair (bp) homology arms flanking a donor gene cassette ***Ghetti et al. (2021)***, whereas the other study incorporated 500-1000 bp homology arms into a donor plasmid ***Philip et al. (2025)***. The short homology arm method can speed up the cell line generation process, as the homology arm can be added to the pair of PCR primers when the donor template is generated, thus eliminating the need for molecular cloning. The difference in the homology arm length may explain the distinct percentage of homozygous knock-in cells between our study and the previous study ***Philip et al. (2025)***, as shown previously ***Li et al. (2014); Zhang et al. (2017); Chai et al. (2024)***. While our approach does not require obtaining homozygous cells, increasing the length of the homology arm by synthesizing longer primers may further improve the efficiency to successfully generate knock-in cells via higher HDR activity. Notably, however, evidence suggests that the optimal stoichiometric ratio of Cas9, guide RNA, and donor dsDNA may play a dominant role in determining the HDR-mediated knock-in efficiency, potentially overriding the effects of homology arm length on the integration efficiency ***Shirk et al. (2026)***. In addition to optimizing these ratios, several other strategies may be considered in future studies to improve the HDR knock-in efficiency. We utilized dsDNA donor templates for the HDR because they are commonly used for the larger inserts and are easy to prepare ***Jin et al. (2025)***, though they often suffer from lower knock-in efficiency ***Renaud et al. (2016)***, higher cellular toxicity ***Shy et al. (2023)***, and the risk of random integration ***Roth et al. (2018)***. In contrast, single-strand DNA (ssDNA) templates often provide higher HDR efficiency ***Renaud et al. (2016)***, lower cellular toxicity ***Shy et al. (2023)***, and lower random integration ***Roth et al. (2018)***, but they are typically limited to the smaller inserts and can be challenging to prepare ***Velangani et al. (2026)***. A notable exception is circular single-stranded DNA (cssDNA), generated by M13 phage, which can reach a size up to 9 kb ***Xie et al. (2025); Ducani et al. (2013); Shepherd et al. (2019)***. CssDNA can be used as a vector to obtain single-stranded templates for HDR. Studies suggest that cssDNA serves as an HDR donor with superior efficiency for CRISPR-based editing compared to linear ssDNA ***Iyer et al. (2022); Letort et al. (2025)***. Even with the optimal stoichiometric ratio of Cas9, guide RNA, and an ideal format of donor template, it is challenging to ensure the donor template is present at the double-strand break (DSB) site. To bridge these gaps, novel strategies have been developed, such as covalently tethering the DNA donor template to the Cas9-guide RNA ribonucleoprotein (RNP) to ensure the presence of the donor DNA at the DSB site with the RNP complex ***Aird et al. (2018); Savic et al. (2018)***. Additionally, the use of CRISPR lipid nanoparticle–spherical nucleic acids (CRISPR LNP–SNAs) for the delivery of CRISPR gene editing components has shown promise in enhancing cellular uptake and improving HDR editing efficiency ***Han et al. (2025)***. Use of these new technologies may further improve the efficacy of our strategy.

### Advantages of our approach

An apparent benefit of our methodology is the ability to bypass single-cell cloning. This not only speeds up the generation of knock-in lines but also mitigates the impact of clone-specific artifacts. The previous study underscored the presence of heterogeneity in supposedly homogenous cell lines ***Westermann et al. (2022)***, and avoiding the selection of clonal populations would increase the reproducibility of biological experiments. Second, the use of heterozygous cells that carry a knock-in cassette as well as the null GOI allele for functional assays may have an additional advantage. One functional copy of the heterozygous cell is often enough to maintain the function of the gene ***Bartha et al. (2018)***, yet the function is more easily disrupted by the degrader drug, as the expression level is already half compared to homozygous cells (as shown in **Figure 4B, 5B, and 6B**). We also addressed the issue in which degron tags themselves decrease the expression levels of fusion proteins by inserting minimal promoters upstream of the GOIs. We showed that minimal promoters can indeed increase fusion protein expression, and that the extent of this increase varies widely across different degron tags and GOIs (**Figure 3**). This suggests that we need to test combinations of tags and mini-promoters to find the optimal expression level of the POI, at which the fusion proteins can maintain physiological function in the absence of the drug, but cannot do so upon degradation by the degrader drug. Given that our method saves time by skipping the single-cell cloning step for creating degron-tag knock-ins, it allows users to test multiple combinations to find the “sweet spot”.

### Limitations and potential modifications for improvement

One disadvantage of our current approach is that it is limited to N-terminal tagging, as we need to knock out the non-knock-in allele. Our strategy is not applicable if the degron tag needs to be fused to the C-terminus of the POI. Secondly, the decreased basal expression level of the degron-tag protein (**Figure 1D-F and 2B-C**) could be due to our bicistronic donor construct that contains a selection marker (blasticidin resistance gene) followed by P2A and degron-tag. The self-cleaving peptide P2A induces ribosomal skipping during translation, creating two proteins from a single open reading frame ***Donnelly et al. (2001)***. It was shown that the expression level of the second gene is lower than the first gene, possibly because of the ribosome fall-off ***Liu et al. (2017); Donnelly et al. (2001)***. Therefore, removing the selection marker and P2A site after successful knock-in via a Cre-loxP system may be an alternative strategy to alleviate the reduced expression of degron-tagged fusion proteins. However, collecting cells in which the selection marker and P2A are cleaved off requires an additional strategy, such as single-cell sorting or fusing the selection marker to GFP to sort GFP-negative cells. The addition of GFP to the knock-in construct may further decrease the homologous recombination efficiency, as the size of the donor DNA increases. Thirdly, although controlling the basal expression level of a degron-fusion protein by inserting a mini-promoter has proven effective (**Figure 3**), the mini-promoter may alter dynamic gene regulation. While the short-EF promoter and minimal CMV promoter that we used are considered core promoters ***Ede et al. (2016); Li et al. (2023)***, binding sites for basal transcription factors (e.g., TFIID) ***Kadonaga (2012); Danino et al. (2015)***, the core promoters can also affect the size of transcriptional bursts, which are primarily regulated by enhancers ***Larsson et al. (2019)***. Thus, gene expression kinetics may be changed through the mini-promoter insertion. Lastly, although we validated this system in near-diploid RPE1-hTERT cells, testing this methodology across a broader range of cell lines remains a priority to demonstrate its robustness across diverse genetic backgrounds.

### Recommended strategy for generating degron-tag knock-in cells

In addition to establishing the heterozygous knock-in degron-tag system, we compared three widely used degron tag systems: the dTAG system, the Halo-PROTAC system, and the IKZF3d system. The results showed that the expression levels and degradation efficiencies of FKBP12^F36V^ and HaloTag are comparable and vary among GOIs, while the expression of the fusion proteins is often lower than the intrinsic expression levels of the GOIs. In contrast, IKZF3d-tagged proteins often exhibit poor expression levels relative to the endogenous protein (**Figure 2, Figure 3, and Figure 3-figure supplement 1 and 2**). Additionally, as mentioned above, inserting a short EF promoter can ameliorate the reduced expression of degron-fusion proteins. With these noted, the recommended strategy for generating heterozygous knock-in cell lines is to make 4 cell lines per GOI, each expressing two different degron tags (FKBP12^F36V^ and HaloTag) under the control of the original or short EF promoter. Each fusion protein likely has a different expression level and degradation efficiency, allowing users to find the best conditions for functional assays.

## Conclusion

By shifting from the single-cell cloning to the heterozygous approach, our method addresses the labor-intensive nature of the traditional approach for generating degron tag knock-in cells. The method also reduces phenotypic variability often associated with clonal isolation and provides a strategy to mitigate the instability of the degron-tag fusion proteins. Thus, our new method expands the applicability and scalability of the valuable degron-tag technology.

## Methods and Materials

### Vector construction

The donor template vectors that served as templates for PCR reactions to generate donor double-stranded DNA (donor dsDNA) were generated in this study and are available from Addgene (#251478, #251479, #251480, #251481, #251482, #251483, #251484, #251485, and #251486). They contain the donor cassette that consists of a mini-promoter (optional), a blasticidin resistance gene, a V5 epitope tag, a specific conditional degron tag, and a rigid linker (EAAAK_3_). The V5-degron tag-rigid linker cassette (Addgene plasmids #185775, #185774, #185772) and blasticidin resistance gene taken from pWPXLd/LAP-C/blast/long EF/DEST ***Kanie et al. (2025)***, mini-promoters taken from pCS2+ vector and pWPXLd/LAPC/blast/short EF/DEST ***Kanie et al. (2025)*** were amplified by polymerase chain reaction (PCR) and assembled into either pUC19 or pENTR221 using Gibson assembly master mix (E2611S, NEB). The insert sequence was validated at Eurofins Genomics by Sanger Sequencing. Sequence and maps for all the vectors are available through Addgene (#87079).

### Cell line and Cell culture

RPE cells stably expressing Blue Fluorescent Protein (BFP)-Cas9 (RPE-BFP-Cas9) ***Kanie et al. (2017)***, and 293T cells were grown in DMEM/F-12 (12400-024, Invitrogen) supplemented with 10% FBS (100-106, Gemini), 1xGlutaMax (35050-061, Gibco), 100 U/mL Penicillin-Streptomycin (15140122, Thermo Fisher Scientific) at 37 °C in 5% CO_2_. Both cell lines were authenticated via a short-tandem-repeat-based test. The authentication was performed by the MTCRO-COBRE Cell Line Authentication Core of the University of Oklahoma Health Science Center. Mycoplasma negativity of the original cell lines (RPE-BFP-Cas9 and 293T) grown in antibiotic-free media was confirmed by a PCR-based test (G238, Applied Biological Materials).

### Compounds

FKBP12 PROTAC dTAG-13 (HY-114421, MedChemExpress), Pomalidomide (HY-10984, MedChemExpress), HaloPROTAC3 (AOB36136, Aobious), were dissolved in dimethyl sulfoxide (DMSO) at a stock concentration of 1 mM and stored in −20°C. Working solutions were prepared by adding stock solution to the culture media and typically contained 0.1% of DMSO.

### gRNA preparation

crRNAs were designed using the ‘Custom Alt-R CRISPR-Cas9 guide RNA’ tool and ordered through IDT. crRNA and tracrRNA (IDT) were resuspended in Nuclease Free Duplex Buffer (IDT) to a final concentration of 100 µM on ice. To prepare guide RNA, 5 µL of 100 µM crRNA and 5 µL of 100 µM tracrRNA were mixed, heated at 95 ^°^C for 5 min, and cooled down to room temperature for 5 min. The annealed gRNA was used for transfection immediately. The crRNA sequence for each GOI is listed in **Table 2**.

**Table 2.**
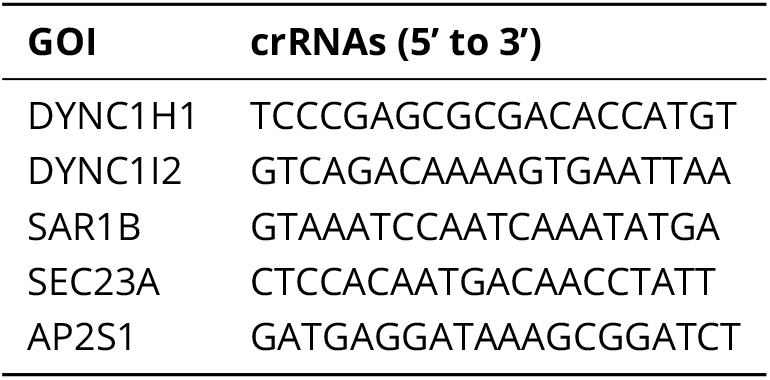
crRNA sequence used in this study.

### Homology-directed repair (HDR) donor double-strand DNA (donor dsDNA) preparation

Donor dsDNAs carrying Left Homology Arm (LHA)-promoter (optional)-blasticidin resistance-P2A-V5-degron-Right Homology Arm (RHA) cassette were generated by PCR using a pair of ∼60 bp oligonucleotides that consists of 40 bp of homology arm and ∼20 bp that anneals with the donor DNA cassette, the donor template vector mentioned above as a template, and Q5 High-Fidelity DNA Polymerase (M0491S, New England BioLabs). The PCR products were separated by DNA electrophoresis and purified using the Zymoclean Gel DNA Recovery Kit (D4002, Zymo Research) according to the manufacturer’s instructions. The concentrations of the recovered Donor dsDNAs were measured using a NanoDrop. The primers used for amplifying each donor dsDNA are listed in **Table 3**. The primer sequence consists of the homology arm (shown in *lowercase, with the optimized codon shown in red*), followed by the Kozak sequence (optional, shown in **Bold**), and the annealing sequence (shown in *uppercase*) to the vectors containing the donor template.

**Table 3.**
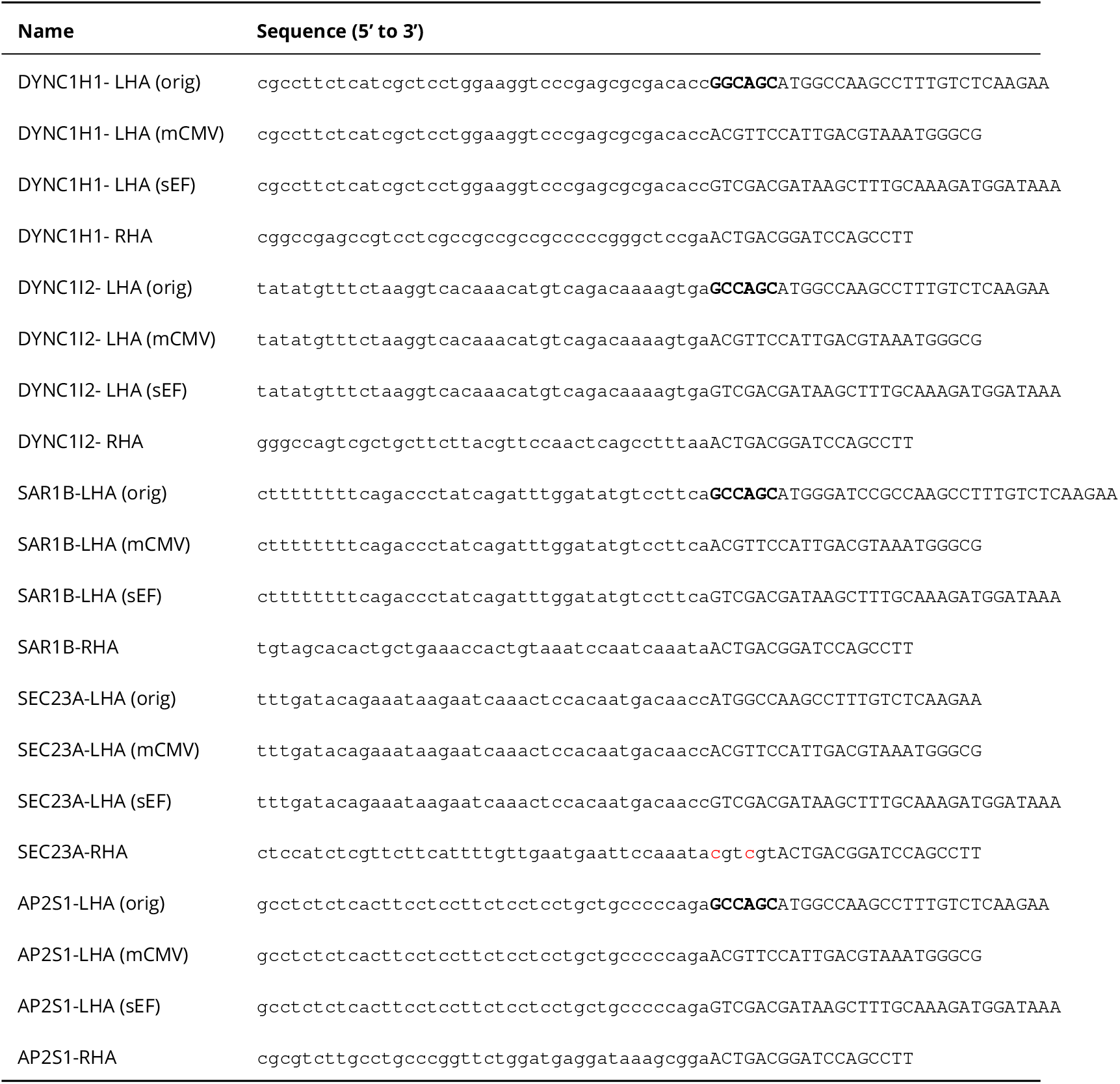
List of the PCR primers for donor dsDNA preparation.

### Generation of stable knock-in cell lines expressing a conditional degron tag

RPE-BFP-Cas9 cells were plated into a 24-well plate (FB012929, Fisherbrand) at a density of 5 × 10^4^ cells per well and grown for 24 hours. Before transfection, two separate mixtures were prepared in a 1.5 ml microcentrifuge tube (05-408-129, Fisher Scientific): Mixture 1 [1.5 μl lipofectamine3000 (L3000001, Invitrogen) in 23.5 μl OPTI-MEM (31985062, Gibco)] and Mixture 2 (30 fmol Donor dsDNA, 15 pmol gRNA, and 1 μl P3000 in 24 μl OPTI-MEM). Both mixtures were incubated at room temperature for 5 minutes. Mixture 2 was then added to Mixture 1, mixed thoroughly by tapping the tube, followed by a brief centrifugation, and was incubated at room temperature for 15 minutes. The transfection mixture was added to the RPE-BFP-Cas9 cells and incubated for 72 hours in 5% CO_2_ incubator at 37°C. The cell lines were then selected with 10 μg/ml blasticidin (14499, Cayman Chemical) until the negative control cells (without infection) reached complete mortality. The selected cells were expanded and subjected to immunoblotting, immunofluorescence, or genomic PCR analyses to determine knock-in/knock-out efficiency.

### Genomic DNA analyses

Genomic DNA used for genomic PCR was prepared by lysing 5 × 10^5^ cells in lysis buffer [10 mM Tris-HCl (pH 7.6), 50 mM NaCl, 6.25 mM MgCl_2_, 0.045% NP40, and 0.45% Tween-20] with Proteinase K (FEREO0491, Thermo Scientific, 200 μg/ml final concentration) treatment for 1 hour at 56 °C. Proteinase K was subsequently inactivated by heating the samples to 95°C for 15 minutes. The resulting lysate was used directly as a template for genomic PCR. Genomic PCR was performed using the template described above, primers listed in **Table 4**, and DreamTaq DNA Polymerase (EP0702, Thermo Scientific). For genotyping shown in **Figure 1B**, the PCR products were run on a 1% agarose gel in 0.5x TAE buffer containing 0.1 μg/ml ethidium bromide at a constant voltage of 100V until the dye front had migrated across approximately two-thirds of the gel length, and the DNA bands were visualized on EpiChemII Darkroom (UVP). For the TIDE analyses, the PCR products were separated by a 1% agarose gel in 0.5x TAE buffer containing MaestroSafe (MR-031203, Transilluminators), and the bands corresponding to the non-knock-in allele were visualized on Blue Light Transilluminator (NEB-SLB-01W, Transilluminator.com) and excised. The PCR products were purified using the Zymoclean Gel DNA Recovery Kit (D4002, Zymo Research) according to the manufacturer’s instructions and Sanger-sequenced by Eurofins Genomics. The sequences were analyzed using the Inference of CRISPR Edits (ICE CRISPR Analysis, 2025, v3.0. EditCo Bio) to determine the percentage of insertions/deletions.

**Table 4.**
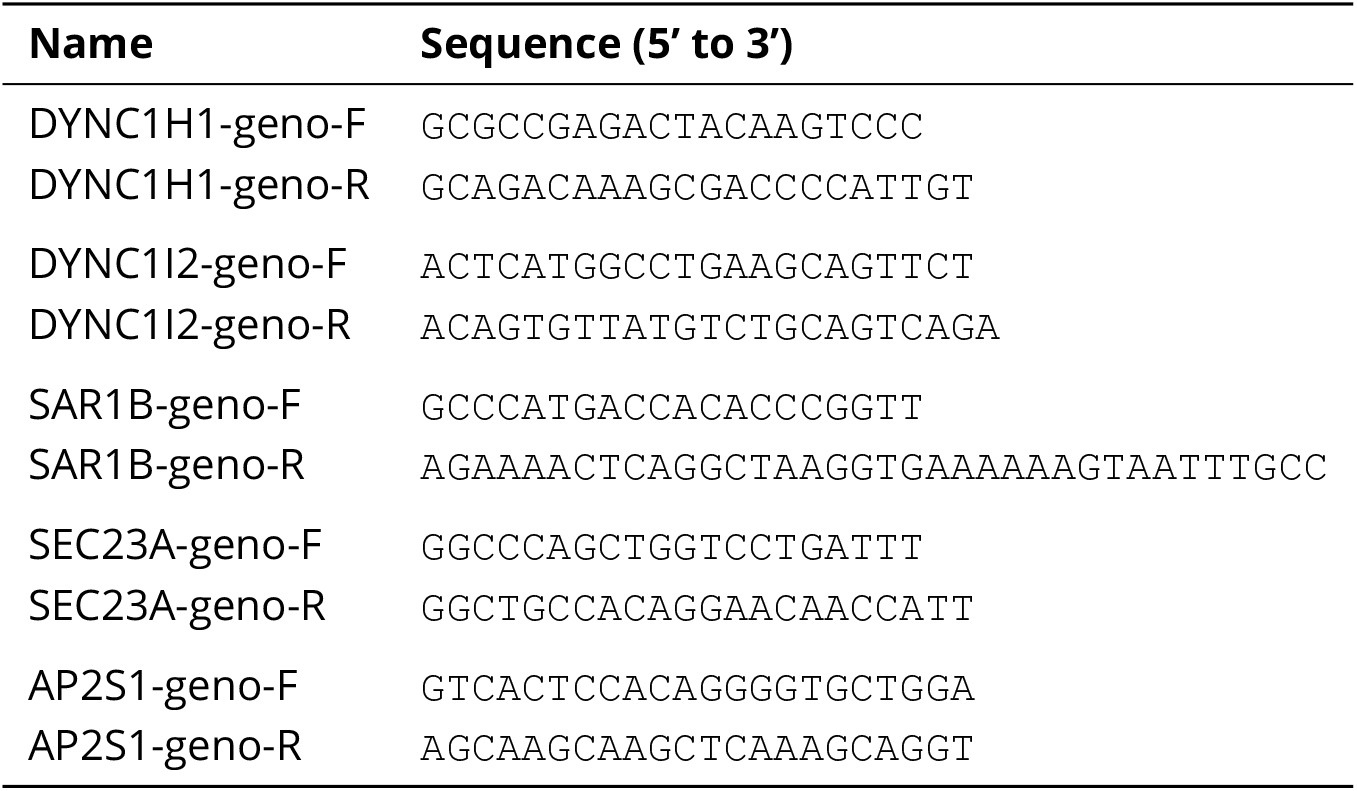
PCR primers for genotyping used in this study.

### Single-Cell Clonal Isolation

Monoclonal cell lines were established from bulk populations of degron tag knock-in cells. Briefly, cells were dissociated using 0.05% Trypsin-EDTA (25300062, Thermo Fisher Scientific) and resuspended in PBS. To remove aggregates, the suspension was passed through a 35 μm nylon mesh. Single-cell sorting was performed on a FACSAria IIIu equipped with a 100 μm nozzle, and doublet depletion was performed using FSC-H vs. FSC-A and SSC-H vs. SSC-A gating. Individual cells were sorted into 96-well plates (12-556-008, Fisher Scientific) containing 100 μL of a 1:1 mixture of fresh DMEM/F12 containing 10% FBS and the conditioned media. The conditioned media were harvested from knock-in bulk cells cultured in DMEM/F-12 containing 10% FBS for 48 hours and filtered through a 0.22 μm PES membrane (431117, Corning).

### Immunoblot

For immunoblotting, cells were grown to confluent in a 6-well plate and lysed in 100 μl of IGEPAL CA-630 lysis buffer {25 mM HEPES-NaOH [pH 7.4], 150 mM NaCl, and 0.3% IGEPAL CA-630 (I8896-100ML, SIGMA)} containing 1% Halt™ Protease Inhibitor Cocktail (87785, Thermo Scientific) followed by clarification of the lysate by centrifugation at 15,000 g at 4°C for 10 min. 72.5 μl of the clarified lysates were then mixed with 25 μl of 4xLaemmli sample buffer (200 mM Tris-HCl [pH 6.8], 8% SDS, 40% glycerol, and 0.02% Bromophenol Blue) and 2.5 μl of 2-mercaptoethanol (M3148, SIGMA), and incubated at 95°C for 5 min. Proteins were separated in 10%, 12%, or top 10%-bottom 12% half-half Bis-Tris protein gels in SDS-PAGE running buffer (25 mM Tris, 192 mM Glycine, 0.1% SDS, pH 8.3), then transferred onto Immobilon™-FL PVDF Transfer Membranes (IPFL00010, MilliporeSigma) in Towbin Buffer (25 mM Tris, 192 mM glycine, pH 8.3). Membranes were incubated in TrueBlack® WB Blocking Buffer (23013A-500ML, Biotium) for 30 min at room temperature and then probed overnight at 4°C with the primary antibody listed in **Table 5**, diluted in TrueBlack® WB Antibody Diluent (23013B-1L, Biotium). Next, membranes were washed 3 x 5 min in TBST buffer (20 mM Tris, 150 mM NaCl, 0.1% Tween 20, pH 7.5) at room temperature, incubated with the secondary antibodies listed in **Table 5** diluted in TrueBlack® WB Antibody Diluent for 30 min at room temperature, then washed 3 x 5 min in TBST buffer. Membranes were scanned on a Sapphire FL biomolecular imager (Azure Biosystems), and proteins were detected at wavelengths 685 and 784 nm. Western blots were quantified by either Fiji or AzureSpot Pro (Azure Biosystems).

**Table 5.** List of antibodies used in this study.

| Antigen | Supplier | Cat# | Dilution |
| --- | --- | --- | --- |
| V5 | Cell Signaling | 13202S | 1:1000 (WB) |
| DYNC1H1 | Proteintech | 12345-1-AP | 1:500 (WB); 1:1000 (IF) |
| DYNC1I2 | Proteintech | 12219-1-AP | 1:1000 (WB) |
| SAR1B | Proteintech | 22292-1-AP | 1:1000 (WB) |
| SEC23A | Abcam | ab179811 | 1:1000 (WB) |
| AP2S1 | Proteintech | 15634-1-AP | 1:1000 (WB) |
| α-Tubulin | Santa Cruz | sc-32293 | 1:500 (WB); 1:1000 (IF) |
| TGN46 | BioRad | AHP500GT | 1:1000 (IF) |
| CF680 Goat Anti-Mouse | Biotium | 20065-1 | 1:20000 (WB) |
| CF770 Goat Anti-Rabbit | Biotium | 20078-1 | 1:20000 (WB) |
| Alexa Fluor 488 Donkey anti-Sheep IgG | Invitrogen | A11015 | 1:2000 (IF) |
| Alexa Fluor 568 Goat Anti-Rabbit IgG | Invitrogen | A11036 | 1:2000 (IF) |
| Alexa Fluor 647 Goat Anti-Mouse IgG1 | Invitrogen | A21240 | 1:2000 (IF) |

### Induction of protein degradation by the degrader drugs for immunoblot analyses

Cells were plated in a 6-well plate (12-556-004, Thermo Scientific) at a density of 2 × 10^5^ cells/well and grown for 48 hr. After the cells were treated with the indicated compound or DMSO for 0-, 1-, 2-, 3-, 6-, 9-, 12-, and 24-hr, all cell lysates were prepared simultaneously for immunoblotting.

### Immunofluorescence

For wide-field microscopy, cells were grown on acid-washed 12 mm #1.5 round coverslips (71861-057, Electron Microscopy Sciences) and fixed in 4% paraformaldehyde (15710, Electron Microscopy Sciences) diluted in phosphate-buffered saline (PBS) for 15 min at room temperature. The primary antibodies used for immunofluorescence are listed in **Table 5**. After permeabilization in immunofluorescence (IF) buffer [3% bovine serum albumin (BP9703100, Fisher Scientific), 0.02% sodium azide (71448-16, Fisher Scientific), and 0.1% IGEPAL® CA-630 in PBS] for 30 min at room temperature, cells were incubated with primary antibody listed in **Table 5** in IF buffer for 3-4 hours at room temperature, followed by rinsing with IF buffer five times. The samples were then incubated with fluorescent dye-labeled secondary antibodies (listed in **Table 5**) in IF buffer for 1 hour at room temperature, followed by rinsing with IF buffer five times. After nuclear staining with 4’,6-diamidino-2-phenylindole (DAPI) (D9542-1MG, SIGMA) in IF buffer at a final concentration of 0.5 μg/ml, coverslips were mounted with Fluoromount-G (0100-01, SouthernBiotech) onto glass slides (22-339-411, Epredia). Images were acquired on a Nikon Ti2 inverted fluorescence microscope equipped with a Fusion BT camera (Model: C15440-20UP, Hamamatsu Photonics), 405 nm, 488 nm, 561 nm, and 642 nm lasers, and CSU-W1 SoRa Confocal Scanner Unit. A 20x NA0.75 plan-apochromat objective lens (MRD00205, Nikon) with the spinning disk confocal was used for mitosis arrest assays. A 60x NA1.4 plan-apochromat Lambda objective lens (MRD01605, Nikon) with a spinning-disk confocal was used for Golgi dispersal and transferrin internalization analyses. A 100x NA1.49 SR HP apochromat TIRF objective lens (MRD01995, Nikon) with the SoRa module was used for super-resolution transferrin internalization analysis. All the raw image data are available through BioImage Archive (Accession ID: S-BIAD2849).

### Mitosis arrest assays and Golgi dispersal assays

Cells were plated into a 6-well plate at a density of 2 × 10^5^ cells/well and grown for 48 hr. After treatment with the indicated compound or DMSO for 3-, 6-, 9-, 12-, 24-hr, all the cells were fixed at the same time in 4% paraformaldehyde diluted in PBS. After permeabilization, the cells used for mitotic arrest assays were stained with DAPI in IF buffer at a final concentration of 0.5 μg/ml for 5 min. The cells used for Golgi dispersal assays were stained with either anti-TGN46 antibody only, or anti-TGN46 antibody combined with anti-DYNC1H1 antibody, and anti-*α*-Tubulin. At least six images from different fields per sample were captured for typical analysis. Typically, roughly 200 cells were analyzed per experiment. The exact number of cells we analyzed in each sample is available in the corresponding Source data in the BioImage Archive (Accession ID: S-BIAD2849). The percentages of mitotically arrested cells and cells with dispersed Golgi were manually counted using the NIS-Elements software (Nikon).

### Transferrin internalization assays

Cells were plated into an 8-well chambered coverslip (80807-90, iBidi) at a density of 5 × 10^4^ cells/well and grown for 48 hours. Cells were treated with HaloPROTAC3 or DMSO for 6 hours and were kept on ice for 10 min. After the cold treatment, the cells were incubated with 25 μg/ml Transferrin Alexa Fluor™ 488 conjugate (T13342, Invitrogen) in complete media without phenol red (11-039-021, Gibco) at 37°C for 15 minutes and washed with PBS 3 times. All the 60x spinning disk confocal live cell images were captured in the CSU-W1 mode to show the inhibition of the transferrin internalization. The 100x SoRa super-resolution Z-stack live cell images were captured in the SoRa mode with the 2.8x magnifier with the 0.175 μm z-increments to ensure Nyquist sampling. Images were then computationally deconvolved using the NIS-Element software.

### Data availability

All data generated or analyzed during this study are included in the manuscript and supporting files; source data files have been provided for all figures. All vectors have been deposited in Addgene (#87079). All immunoblot and immunofluorescent data have been deposited at BioImage Archive (Accession ID: S-BIAD2849).

## Acknowledgments

We would like to thank the members of the Kanie lab for their insightful discussions. The Nikon CSU-W1/SoRa super-resolution spinning disk confocal microscope is supported by an Equipment Grant from the Presbyterian Health Foundation (PHF) and the OUHC Department of Cell Biology. The cell authentication service performed by MTCRO-COBRE Cell line authentication core of the University of Oklahoma Health Campus was supported partly by P20GM103639 and National Cancer Institute Grant P30CA225520 of the National Institutes of Health (NIH). The single-cell sorting service was performed by the Flow Cytometry Core of the Oklahoma Medical Research Foundation. This project was supported in part by the National Institute of General Medical Sciences (NIGMS, Cellular and Molecular Geroscience CoBRE: P20GM12558) awarded to the University of Oklahoma, NIH grants P20GM103447, 1R35GM151013, and PHF research support grant to TK.

## Author Contributions

**Beibei Liu**: Conceptualization, Data curation, Formal analysis, Validation, Investigation, Visualization, Methodology, Writing–original draft, Writing–review and editing.

**Chao Qi**: Data curation, Formal analysis, Validation, Methodology, Writing–review and editing.

**Tomoharu Kanie**: Conceptualization, Methodology, Validation, Resources, Supervision, Funding acquisition, Project administration, Writing–original draft, Writing–review and editing.

---

**Figure 1—figure supplement 1.**
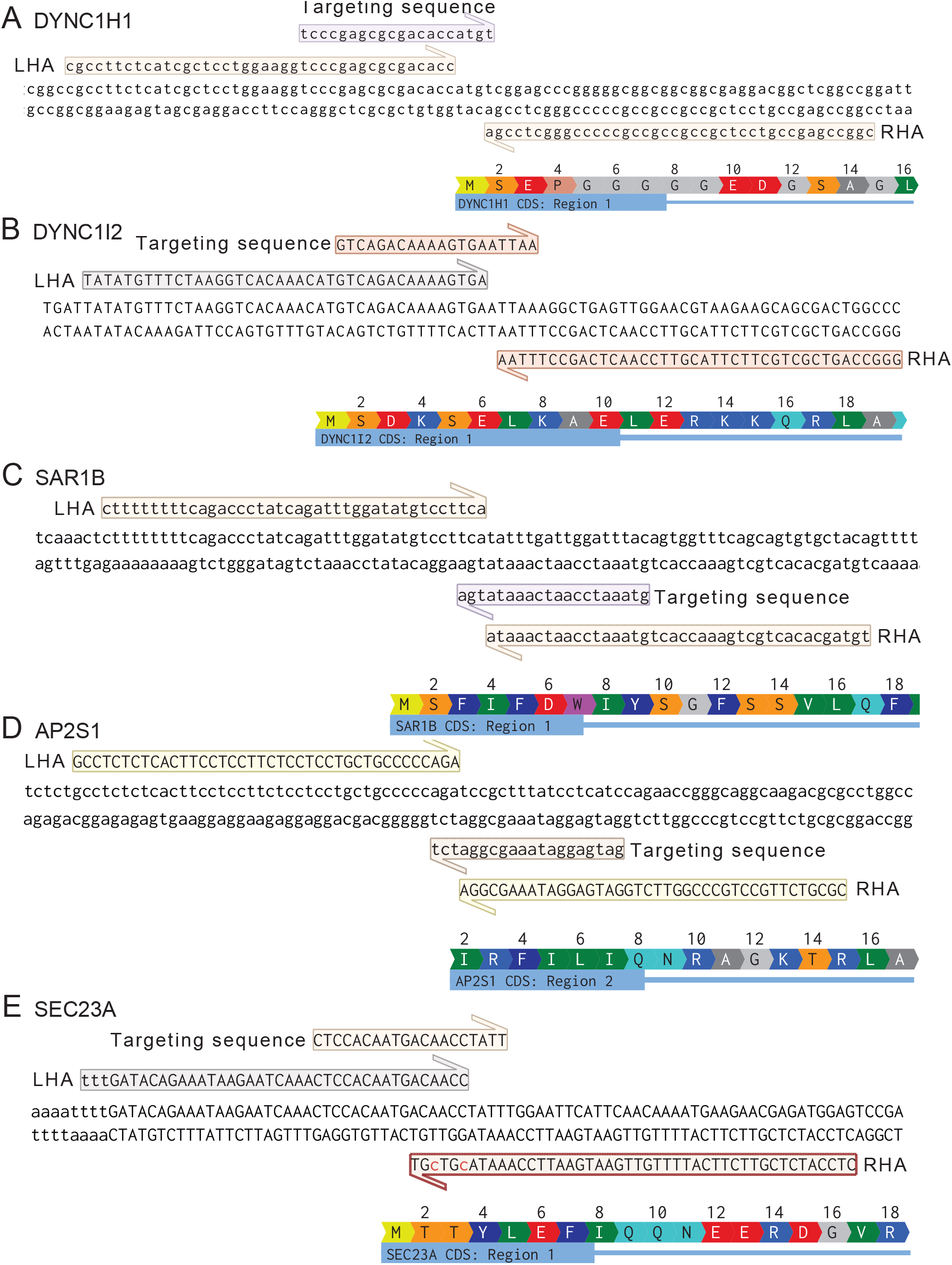
The genomic DNA sequence flanking the CRISPR-Cas9 targeting region for each gene of interest. **(A-E)**. The CRISPR-Cas9 targeting genomic DNA sequence, left/right homology arm (LHA/RHA) sequences used in knock-in cell generation, and partial Coding DNA Sequence (CDS) for human DYNC1H1 **(A)**, DYNC1I2 **(B)**, SAR1B **(C)**, AP2S1 **(D)**, and SEC23A **(E)**. The half-arrows indicate 5’ to 3’ direction of the sequences.

**Figure 1—figure supplement 2.**
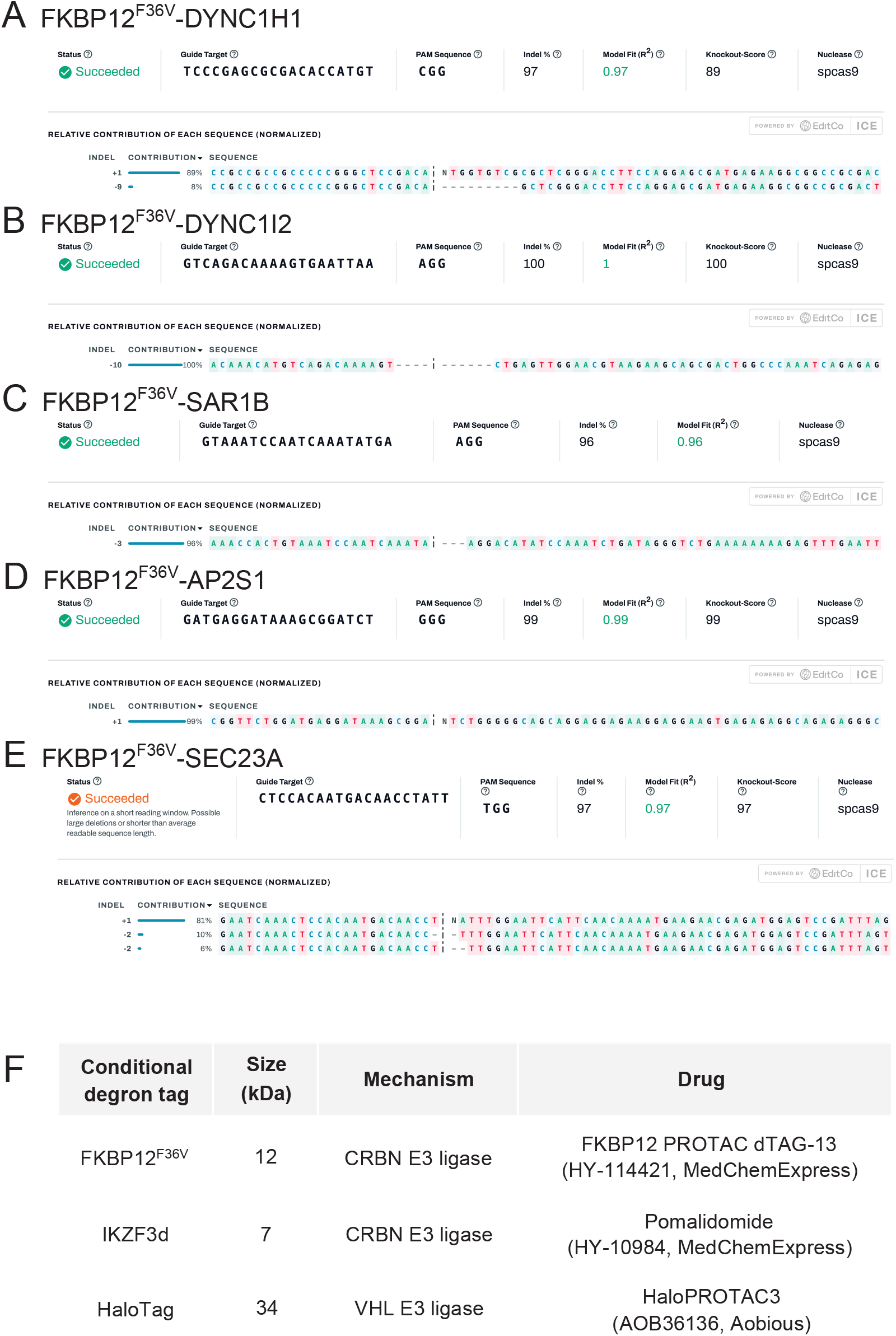
Genomic DNA sequences of the non-knock-in allele in the heterozygous knock-in cells. **(A-E)**. The ICE CRISPR Analysis data showing insertion/deletion harbored in the non-knock-in allele of the heterozygous knock-in RPE-BFP-Cas9 cells expressing FKBP12^*F* 36*V*^ tagged DYNC1H1**(A)**, DYNC1I2 **(B)**, SAR1B **(C)**, AP2S1 **(D)**, and SEC23A **(E)**. The percentage of insertion/deletion was analyzed by the ICE CRISPR analysis tool following the purification of PCR-amplified genomic DNA. Indel % shows the percentage of sequences that are different from control samples; Model Fit (R^2^) shows how well the model fits the data; Knockout Score indicates the percentage of cells with either frameshift or +21bp insertion/deletion. Notably, Figure A shows that 8% of sequences harbor a 9-bp deletion that disrupts the start codon of DYNC1H1. **(F)**. A table summarizing the size, an E3 ubiquitin ligase that is used for the degradation (mechanism), and a drug used in this study for each conditional degron tag.

**Figure 2—figure supplement 1.**
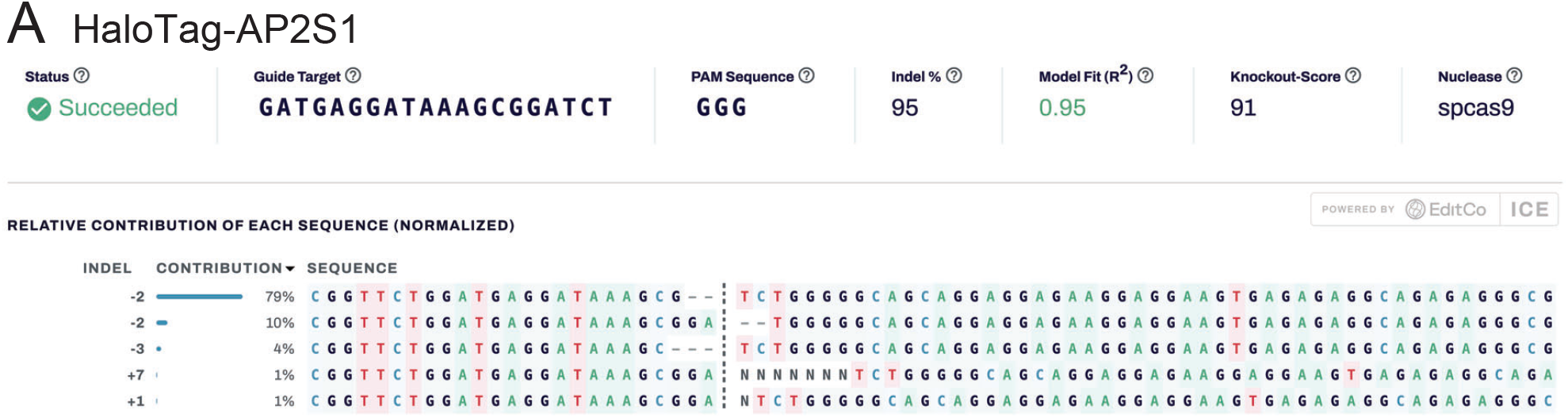
Genomic DNA sequences of the non-knock-in allele in the heterozygous HaloTag-AP2S1 knock-in cells. **(A-E)**. The ICE CRISPR Analysis data showing insertion/deletion harbored in the non-knock-in allele of the heterozygous knock-in RPE-BFP-Cas9 cells expressing AP2S1 fused to HaloTag **(Figure 2C)**. The percentage of insertion/deletion was analyzed by the ICE CRISPR analysis tool following the purification of PCR-amplified genomic DNA. Indel % shows the percentage of sequences that are different from control samples; Model Fit (R^2^) shows how well the model fits the data; Knockout Score indicates the percentage of cells with either frameshift or +21bp insertion/deletion. While the upper-shifted form of AP2S1 compared to the endogenous AP2S1 was observed in the immunoblot analysis (the blue asterisk in **Figure 2C**), the cells have dominant −2 deletions.

**Figure 3—figure supplement 1.**
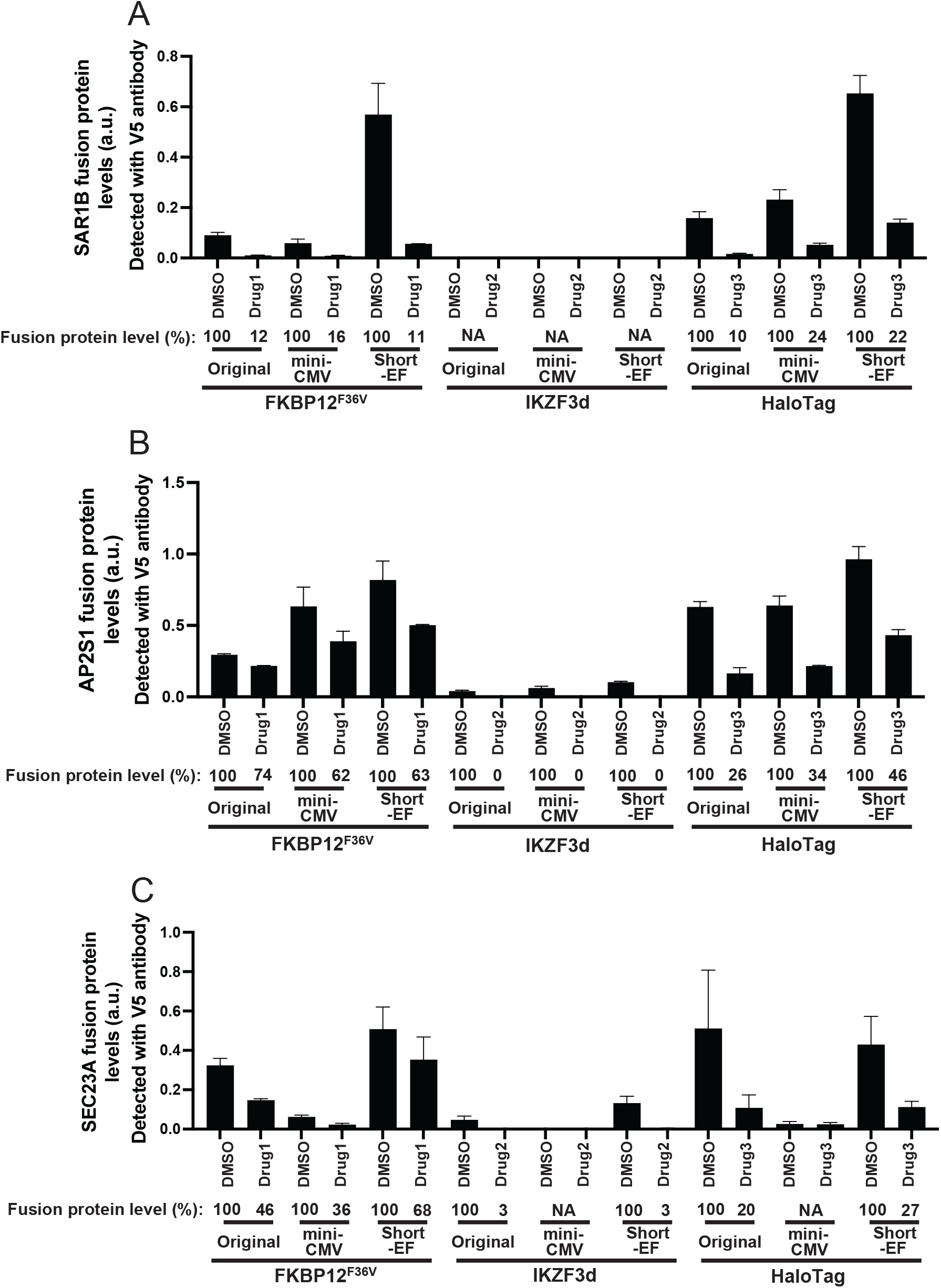
Quantification of the expression levels of the fusion proteins shown in Figure 3. **(A-C)**. Quantification of protein expression in the cells shown in **Figure 3 B-D**. The bands shown in the Immunoblot analysis using the V5-tag antibody were quantified. Fusion protein level (%) was calculated by normalizing the expression level of the drug-treated cells to the DMSO-treated cells. The data is combined from two independent experiments. Error bars represent Mean ± SEM

**Figure 3—figure supplement 2.**
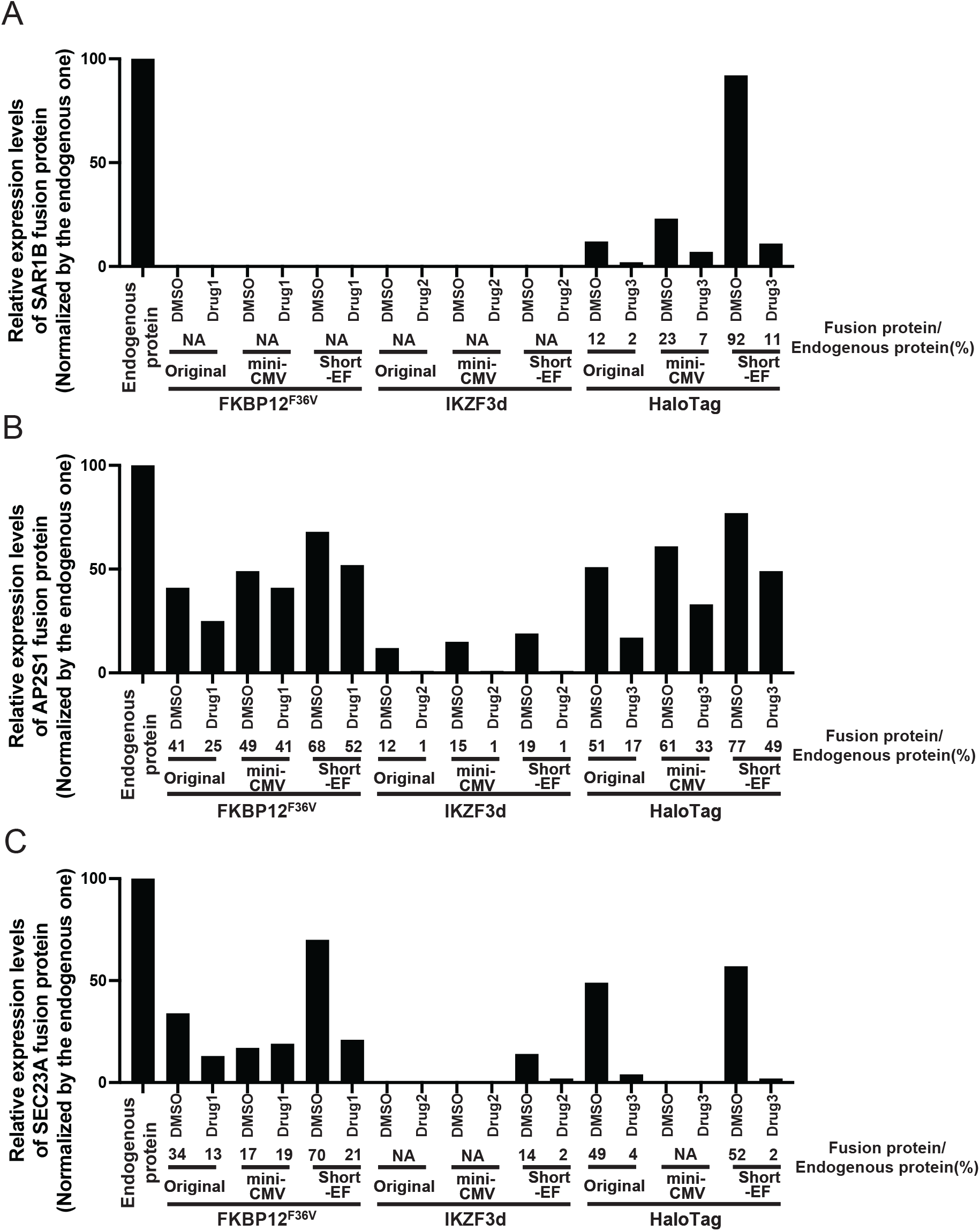
Quantification of the expression levels of the fusion proteins compared with those of the endogenous one. **(A-C)**. Quantification of protein expression in the cells shown in **Figure 3B-D**. The bands shown in the Immunoblot analysis using the SAR1B antibody **(A)**, AP2S1 antibody **(B)**, and SEC23A antibody **(C)** were quantified. Fusion protein level (%) was calculated by normalizing the expression level of the DMSO or drug-treated cells to the respective endogenous protein.

**Figure 3—figure supplement 3.**
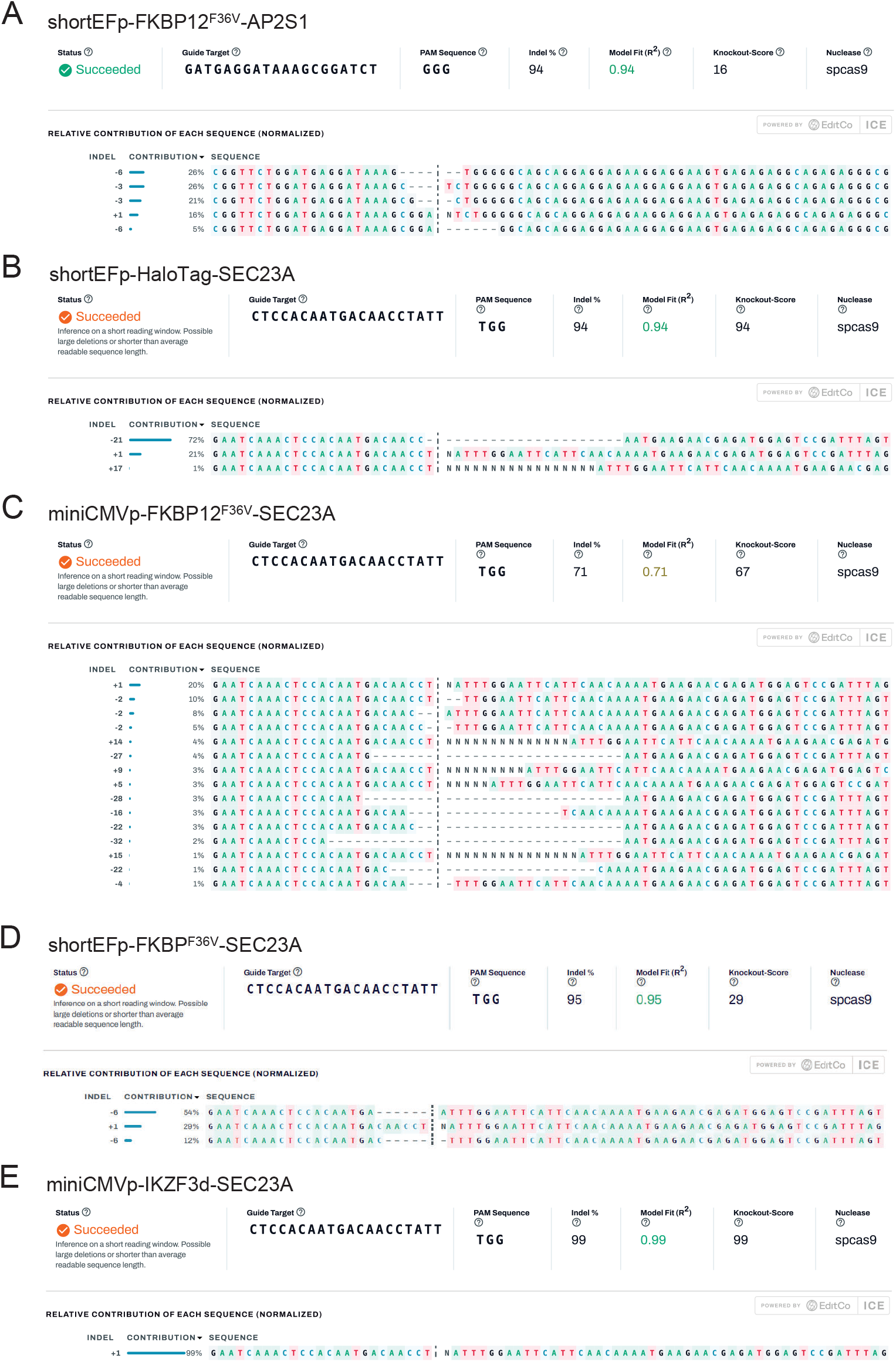
The ICE CRISPR Analysis data of the mini-promoter-regulated degron-tag knock-in cells. **(A-D)**. The ICE CRISPR Analysis data showing insertion/deletion harbored in the non-knock-in allele of the heterozygous knock-in RPE-BFP-Cas9 cells expressing FKBP12F36V-AP2S1 under the control of short-EF promoter **(A)**, HaloTag-SEC23A under the control of short-EF promoter **(B)**, FKBP12F36V-SEC23A under the control of mini-CMV **(C)**, FKBP12^*F* 36*V*^-SEC23A under the control of short-EF promoter **(D)**, and IKZF3d-SEC23A under the control of mini-CMV **(E)**. The percentage of insertion/deletion was analyzed by the ICE CRISPR analysis tool following the purification of PCR-amplified genomic DNA. Indel % shows the percentage of sequences that are different from control samples; Model Fit (R^2^) shows how well the model fits the data; Knockout Score indicates the percentage of cells with either frameshift or +21bp insertion/deletion. Notably, the weak bands detected in the cells expressing IKZF3d-SEC23A under the control of a minimal CMV promoter **(Figure 3D)** cannot readily explain the dominant +1 insertion detected in **E**. A possible explanation for this is that most of the cells may have a huge deletion that prevented the primers from binding, thus the actual deletion was not reflected in the ICE analysis.

**Figure 4—figure supplement 1.**
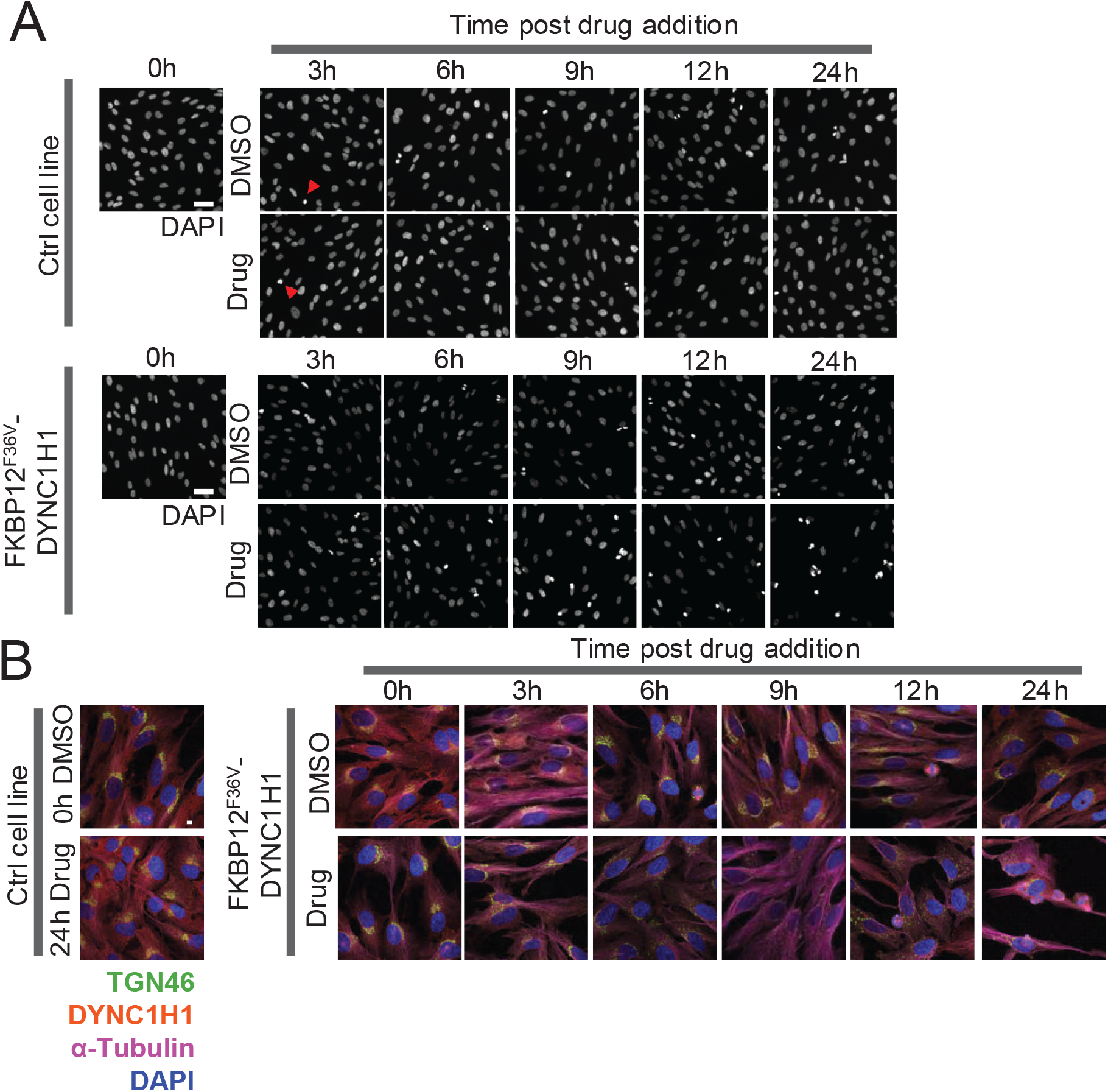
The control experiments for the mitosis arrest assay and the Golgi dispersal assay shown in Figure 4. **(A)**. Mitosis arrest assay in control (the parental RPE-BFP-Cas9 cells) and the heterozygous knock-in RPE-BFP-Cas9 cells expressing FKBP12^*F* 36*V*^-DYNC1H1 cells. The cells were stained with DAPI. Representative images of control (DMSO) or 1 μM dTAG-13 (drug)-treated cells at the indicated time point are shown. The red arrowheads indicate the mitotic cells. Scale bar: 50 μm **(B)**. Golgi dispersal assay in control (the parental RPE-BFP-Cas9 cells) and the heterozygous knock-in cells expressing FKBP12^*F* 36*V*^-DYNC1H1. The cells were stained with the indicated antibodies. Representative images of control (DMSO) or 1 μM dTAG-13 (drug)-treated cells at the indicated time point are shown. Scale bar: 10 μm

**Figure 6—figure supplement 1.**
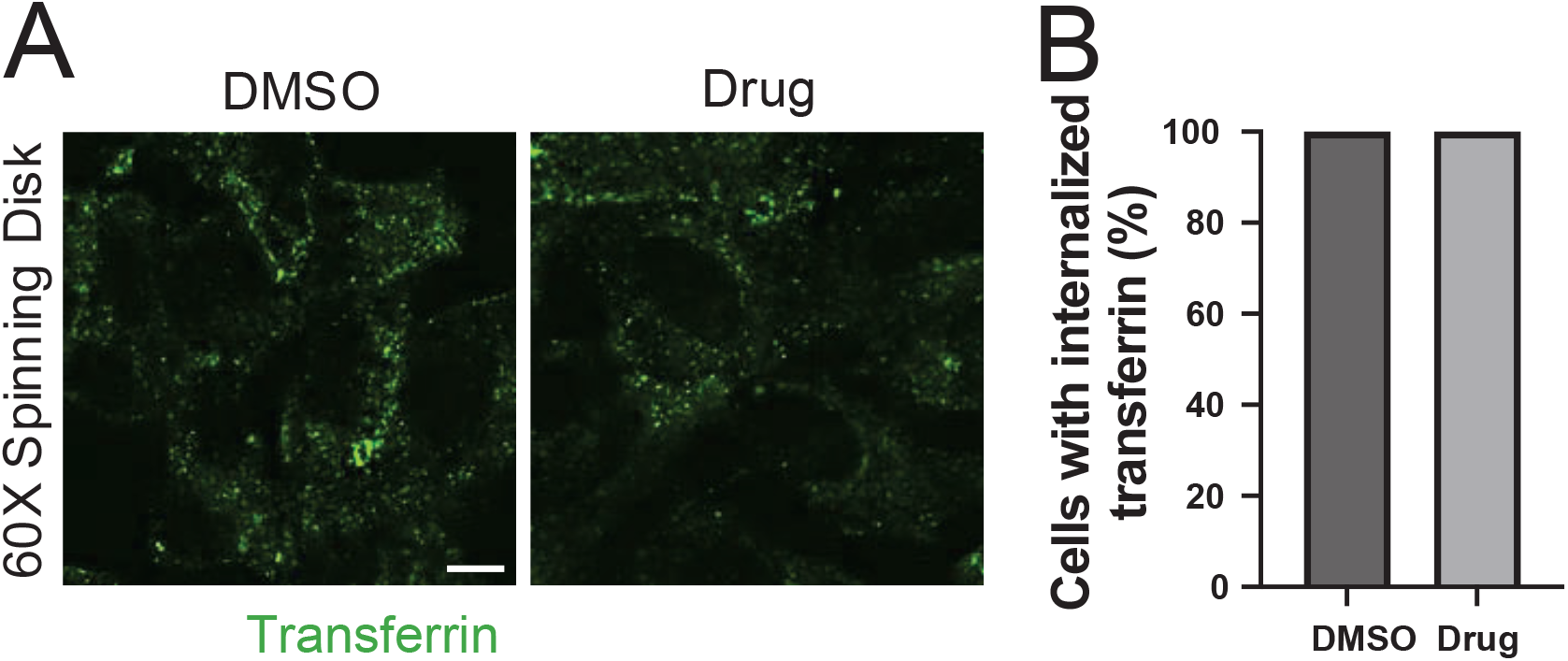
Transferrin internalization assays in the heterozygous knock-in cells expressing FKBP12^*F* 36*V*^-AP2S1 under the control of short-EF promoter. **(A)**. Transferrin internalization assays in the heterozygous knock-in RPE-BFP-Cas9 cells expressing short EF promoter-regulated FKBP12^*F* 36*V*^-AP2S1. The cells were treated with either dimethyl sulfoxide (DMSO) or 1 μM HaloPROTAC3 (drug) for 6 hours and incubated with transferrin-Alexa Fluor 488. Live-cell imaging was performed on a spinning disk confocal microscope. The individual image is from a representative z-slice. Scale bar: 10 μm **(B)**. Quantification of the percentage of the cells with internalized transferrin shown in **Figure 6-figure supplement 1A**. The data is from a representative experiment.

## References

Abuhashem A, Hadjantonakis AK. Generation of knock-in degron tags for endogenous proteins in mice using the dTAG system. STAR Protoc. 2022; 3(3):101660. https://www.ncbi.nlm.nih.gov/pubmed/36097386, doi: 10.1016/j.xpro.2022.101660, abuhashem, Abderhman Hadjantonakis, Anna-Katerina eng R01 HD094868/HD/NICHD NIH HHS/R01 DK127821/DK/NIDDK NIH HHS/R01 HD086478/HD/NICHD NIH HHS/T32 GM007739/GM/NIGMS NIH HHS/P30 CA008748/CA/NCI NIH HHS/F30 HD103398/HD/NICHD NIH HHS/Research Support, N.I.H., Extramural 2022/09/14 STAR Protoc. 2022 Sep 16;3(3):101660. doi: 10.1016/j.xpro.2022.101660. Epub 2022 Sep 7.

Aird EJ, Lovendahl KN, St Martin A, Harris RS, Gordon WR. Increasing Cas9-mediated homology-directed repair efficiency through covalent tethering of DNA repair template. Commun Biol. 2018; 1:54. https://www.ncbi.nlm.nih.gov/pubmed/30271937, doi: 10.1038/s42003-018-0054-2, aird, Eric J Lovendahl, Klaus N St Martin, Amber Harris, Reuben S Gordon, Wendy R eng R01 GM118000/GM/NIGMS NIH HHS/R35 GM119483/GM/NIGMS NIH HHS/R37 AI064046/AI/NIAID NIH HHS/T32 GM008347/GM/NIGMS NIH HHS/England 2018/10/03 Commun Biol. 2018 May 31;1:54. doi: 10.1038/s42003-018-0054-2. eCollection 2018.

Azuma T, Kei T. Super-resolution spinning-disk confocal microscopy using optical photon reassignment. Opt Express. 2015; 23(11):15003–11. https://www.ncbi.nlm.nih.gov/pubmed/26072856, doi: 10.1364/OE.23.015003, azuma, Takuya Kei, Takayuki eng 2015/06/16 Opt Express. 2015 Jun 1;23(11):15003–11. doi: 10.1364/OE.23.015003.

Bartha I, di Iulio J, Venter JC, Telenti A. Human gene essentiality. Nat Rev Genet. 2018; 19(1):51–62. https://www.ncbi.nlm.nih.gov/pubmed/29082913, doi: 10.1038/nrg.2017.75, bartha, Istvan di Iulio, Julia Venter, J Craig Telenti, Amalio eng Review England 2017/10/31 Nat Rev Genet. 2018 Jan;19(1):51–62. doi: 10.1038/nrg.2017.75. Epub 2017 Oct 30.

Bondeson DP, Mullin-Bernstein Z, Oliver S, Skipper TA, Atack TC, Bick N, Ching M, Guirguis AA, Kwon J, Langan C, Millson D, Paolella BR, Tran K, Wie SJ, Vazquez F, Tothova Z, Golub TR, Sellers WR, Ianari A. Systematic profiling of conditional degron tag technologies for target validation studies. Nat Commun. 2022; 13(1):5495. https://www.ncbi.nlm.nih.gov/pubmed/36127368, doi: 10.1038/s41467-022-33246-4, bondeson, Daniel P Mullin-Bernstein, Zachary Oliver, Sydney Skipper, Thomas A Atack, Thomas C Bick, Nolan Ching, Meilani Guirguis, Andrew A Kwon, Jason Langan, Carly Millson, Dylan Paolella, Brenton R Tran, Kevin Wie, Sarah J Vazquez, Francisca Tothova, Zuzana Golub, Todd R Sellers, William R Ianari, Alessandra eng K00 CA212229/CA/NCI NIH HHS/R01 CA233626/CA/NCI NIH HHS/R35 CA242457/CA/NCI NIH HHS/Research Support, N.I.H., Extramural Research Support, Non-U.S. Gov’t England 2022/09/21 Nat Commun. 2022 Sep 20;13(1):5495. doi: 10.1038/s41467-022-33246-4.

Boshart M, Weber F, Jahn G, Dorsch-Hasler K, Fleckenstein B, Schaffner W. A very strong enhancer is located upstream of an immediate early gene of human cytomegalovirus. Cell. 1985; 41(2):521–30. https://www.ncbi.nlm.nih.gov/pubmed/2985280, doi: 10.1016/s0092-8674(85)80025-8, boshart, M Weber, F Jahn, G Dorsch-Hasler, K Fleckenstein, B Schaffner, W eng Research Support, Non-U.S. Gov’t 1985/06/01 Cell. 1985 Jun;41(2):521–30. doi: 10.1016/s0092-8674(85)80025-8.

Brinkman EK, Chen T, Amendola M, van Steensel B. Easy quantitative assessment of genome editing by sequence trace decomposition. Nucleic Acids Res. 2014; 42(22):e168. doi: 10.1093/nar/gku936, 1362–4962 Brinkman, Eva K Chen, Tao Amendola, Mario van Steensel, Bas 293662/ERC_/European Research Council/International Journal Article Research Support, Non-U.S. Gov’t Nucleic Acids Res. 2014 Dec 16;42(22):e168. doi: 10.1093/nar/gku936. Epub 2014 Oct 9.

Buckley DL, Raina K, Darricarrere N, Hines J, Gustafson JL, Smith IE, Miah AH, Harling JD, Crews CM. HaloPROTACS: Use of Small Molecule PROTACs to Induce Degradation of HaloTag Fusion Proteins. ACS Chem Biol. 2015; 10(8):1831–7. https://www.ncbi.nlm.nih.gov/pubmed/26070106, doi: 10.1021/acschembio.5b00442, buckley, Dennis L Raina, Kanak Darricarrere, Nicole Hines, John Gustafson, Jeffrey L Smith, Ian E Miah, Afjal H Harling, John D Crews, Craig M eng T32 GM067543/GM/NIGMS NIH HHS/R01 AI084140/AI/NIAID NIH HHS/F32GM10052101/GM/NIGMS NIH HHS/R01AI084140/AI/NIAID NIH HHS/F32 GM100521/GM/NIGMS NIH HHS/T32GM067543/GM/NIGMS NIH HHS/Research Support, N.I.H., Extramural Research Support, U.S. Gov’t, Non-P.H.S. 2015/06/13 ACS Chem Biol. 2015 Aug 21;10(8):1831–7. doi: 10.1021/acschembio.5b00442. Epub 2015 Jun 23.

Chai R, Guo J, Geng Y, Huang S, Wang H, Yao X, Li T, Qiu L. The Influence of Homologous Arm Length on Homologous Recombination Gene Editing Efficiency Mediated by SSB/CRISPR-Cas9 in Escherichia coli. Microorganisms. 2024; 12(6). https://www.ncbi.nlm.nih.gov/pubmed/38930484, doi: 10.3390/microorganisms12061102, chai, Ran Guo, Jiaxiang Geng, Yue Huang, Shuai Wang, Haifeng Yao, Xinding Li, Tao Qiu, Liyou eng 32200089/National Natural Science Foundation of China/221100320200/Major Special Science and Technology Project of Henan Province/231111320200/Key Research and Development Special Project of Henan Province/242102310336/Science and Technology Development Plan of Henan Province/Switzerland 2024/06/27 Microorganisms. 2024 May 29;12(6):1102. doi: 10.3390/microorganisms12061102.

Charpentier M, Khedher AHY, Menoret S, Brion A, Lamribet K, Dardillac E, Boix C, Perrouault L, Tesson L, Geny S, De Cian A, Itier JM, Anegon I, Lopez B, Giovannangeli C, Concordet JP. CtIP fusion to Cas9 enhances transgene integration by homology-dependent repair. Nat Commun. 2018; 9(1):1133. https://www.ncbi.nlm.nih.gov/pubmed/29556040, doi: 10.1038/s41467-018-03475-7, charpentier, M Khedher, AYY Menoret, S Brion, A Lamribet, K Dardillac, E Boix, C Perrouault, L Tesson, L Geny, S De Cian, A Itier, M M Anegon, I Lopez, B Giovannangeli, C Concordet, JP eng Research Support, Non-U.S. Gov’t England 2018/03/21 Nat Commun. 2018 Mar 19;9(1):1133. doi: 10.1038/s41467-018-03475-7.

Chenouard V, Leray I, Tesson L, Remy S, Allan A, Archer D, Caulder A, Fortun A, Bernardeau K, Cherifi Y, Teboul L, David L, Anegon I. Excess of guide RNA reduces knockin efficiency and drastically increases on-target large deletions. iScience. 2023; 26(4):106399. https://www.ncbi.nlm.nih.gov/pubmed/37034986, doi: 10.1016/j.isci.2023.106399, chenouard, Vanessa Leray, Isabelle Tesson, Laurent Remy, Severine Allan, Alasdair Archer, Daniel Caulder, Adam Fortun, Agnes Bernardeau, Karine Cherifi, Yacine Teboul, Lydia David, Laurent Anegon, Ignacio eng MC_UP_2201/3/MRC_/Medical Research Council/United Kingdom 2023/04/11 iScience. 2023 Mar 14;26(4):106399. doi: 10.1016/j.isci.2023.106399. eCollection 2023 Apr 21.

Cho NH, Cheveralls KC, Brunner AD, Kim K, Michaelis AC, Raghavan P, Kobayashi H, Savy L, Li JY, Canaj H, Kim JYS, Stewart EM, Gnann C, McCarthy F, Cabrera JP, Brunetti RM, Chhun BB, Dingle G, Hein MY, Huang B, et al. OpenCell: Endogenous tagging for the cartography of human cellular organization. Science. 2022; 375(6585):eabi6983. https://www.ncbi.nlm.nih.gov/pubmed/35271311, doi: 10.1126/science.abi6983, cho, Nathan H Cheveralls, Keith C Brunner, Andreas-David Kim, Kibeom Michaelis, Andre C Raghavan, Preethi Kobayashi, Hirofumi Savy, Laura Li, Jason Y Canaj, Hera Kim, JamesS S Stewart, Edna M Gnann, Christian McCarthy, Frank Cabrera, Joana P Brunetti, Rachel M Chhun, Bryant B Dingle, Greg Hein, Marco Y Huang, Bo Mehta, Shalin B Weissman, Jonathan S Gomez-Sjoberg, Rafael Itzhak, Daniel N Royer, Loic A Mann, Matthias Leonetti, Manuel D eng F31 HL143882/HL/NHLBI NIH HHS/R01 GM131641/GM/NIGMS NIH HHS/RM1 HG009490/HG/NHGRI NIH HHS/HHMI/Howard Hughes Medical Institute/Research Support, N.I.H., Extramural Research Support, Non-U.S. Gov’t 2022/03/11 Science. 2022 Mar 11;375(6585):eabi6983. doi: 10.1126/science.abi6983. Epub 2022 Mar 11.

Cloarec-Ung FM, Beaulieu J, Suthananthan A, Lehnertz B, Sauvageau G, Sheppard HM, Knapp D. Near-perfect precise on-target editing of human hematopoietic stem and progenitor cells. Elife. 2024; 12. https://www.ncbi.nlm.nih.gov/pubmed/38829685, doi: 10.7554/eLife.91288, cloarec-Ung, Fanny-Mei Beaulieu, Jamie Suthananthan, Arunan Lehnertz, Bernhard Sauvageau, Guy Sheppard, Hilary M Knapp, David J H F eng 944254/Cancer Research Society/TFRI 1118/Terry Fox Research Institute/283502/Fonds de Recherche du Quebec - Sante/England 2024/06/03 Elife. 2024 Jun 3;12:RP91288. doi: 10.7554/eLife.91288.

Collins BM, McCoy AJ, Kent HM, Evans PR, Owen DJ. Molecular architecture and functional model of the endocytic AP2 complex. Cell. 2002; 109(4):523–35. https://www.ncbi.nlm.nih.gov/pubmed/12086608, doi: 10.1016/s0092-8674(02)00735-3, collins, Brett M McCoy, Airlie J Kent, Helen M Evans, Philip R Owen, David J eng Research Support, Non-U.S. Gov’t 2002/06/28 Cell. 2002 May 17;109(4):523–35. doi: 10.1016/s0092-8674(02)00735-3.

Cullot G, Aird EJ, Schlapansky MF, Yeh CD, van de Venn L, Vykhlyantseva I, Kreutzer S, Mailander D, Lewkow B, Klermund J, Montellese C, Biserni M, Aeschimann F, Vonarburg C, Gehart H, Cathomen T, Corn JE. Genome editing with the HDR-enhancing DNA-PKcs inhibitor AZD7648 causes large-scale genomic alterations. Nat Biotechnol. 2024; https://www.ncbi.nlm.nih.gov/pubmed/39604565, doi: 10.1038/s41587-024-02488-6, cullot, Gregoire Aird, Eric J Schlapansky, Moritz F Yeh, Charles D van de Venn, Lilly Vykhlyantseva, Iryna Kreutzer, Susanne Mailander, Dominic Lewkow, Bohdan Klermund, Julia Montellese, Christian Biserni, Martina Aeschimann, Florian Vonarburg, Cedric Gehart, Helmuth Cathomen, Toni Corn, Jacob E eng 855741-DDREAMM-ERC-2019-SyG/EC | Horizon 2020 Framework Programme (EU Framework Programme for Research and Innovation H2020)/DFG CA 311/4-1/Deutsche Forschungsgemeinschaft (German Research Foundation)/2024/11/28 Nat Biotechnol. 2024 Nov 27. doi: 10.1038/s41587-024-02488-6.

Damhofer H, Radzisheuskaya A, Helin K. Generation of locus-specific degradable tag knock-ins in mouse and human cell lines. STAR Protoc. 2021; 2(2):100575. https://www.ncbi.nlm.nih.gov/pubmed/34151298, doi: 10.1016/j.xpro.2021.100575, damhofer, Helene Radzisheuskaya, Aliaksandra Helin, Kristian eng P30 CA008748/CA/NCI NIH HHS/Research Support, N.I.H., Extramural Research Support, Non-U.S. Gov’t 2021/06/22 STAR Protoc. 2021 Jun 2;2(2):100575. doi: 10.1016/j.xpro.2021.100575. eCollection 2021 Jun 18.

Danino YM, Even D, Ideses D, Juven-Gershon T. The core promoter: At the heart of gene expression. Biochim Biophys Acta. 2015; 1849(8):1116–31. https://www.ncbi.nlm.nih.gov/pubmed/25934543, doi: 10.1016/j.bbagrm.2015.04.003, danino, Yehuda M Even, Dan Ideses, Diana Juven-Gershon, Tamar eng Review Netherlands 2015/05/03 Biochim Biophys Acta. 2015 Aug;1849(8):1116–31. doi: 10.1016/j.bbagrm.2015.04.003. Epub 2015 Apr 28.

Donnelly MLL, Luke G, Mehrotra A, Li X, Hughes LE, Gani D, Ryan MD. Analysis of the aphthovirus 2A/2B polyprotein ‘cleavage’ mechanism indicates not a proteolytic reaction, but a novel translational effect: a putative ribosomal ‘skip’. J Gen Virol. 2001; 82(Pt 5):1013–1025. https://www.ncbi.nlm.nih.gov/pubmed/11297676, doi: 10.1099/0022-1317-82-5-1013, donnelly, MichelleL L Luke, Garry Mehrotra, Amit Li, Xuejun Hughes, Lorraine E Gani, David Ryan, Martin D eng Research Support, Non-U.S. Gov’t England 2001/04/12 J Gen Virol. 2001 May;82(Pt 5):1013–1025. doi: 10.1099/0022-1317-82-5-1013.

Ducani C, Kaul C, Moche M, Shih WM, Hogberg B. Enzymatic production of ‘monoclonal stoichiometric’ single-stranded DNA oligonucleotides. Nat Methods. 2013; 10(7):647–52. https://www.ncbi.nlm.nih.gov/pubmed/23727986, doi: 10.1038/nmeth.2503, ducani, Cosimo Kaul, Corinna Moche, Martin Shih, William M Hogberg, Bjorn eng DP2 OD004641/OD/NIH HHS/Research Support, Non-U.S. Gov’t 2013/06/04 Nat Methods. 2013 Jul;10(7):647–52. doi: 10.1038/nmeth.2503. Epub 2013 Jun 2.

Ede C, Chen X, Lin MY, Chen YY. Quantitative Analyses of Core Promoters Enable Precise Engineering of Regulated Gene Expression in Mammalian Cells. ACS Synth Biol. 2016; 5(5):395–404. https://www.ncbi.nlm.nih.gov/pubmed/26883397, doi: 10.1021/acssynbio.5b00266, ede, Christopher Chen, Ximin Lin, Meng-Yin Chen, Yvonne Y eng DP5 OD012133/OD/NIH HHS/Research Support, N.I.H., Extramural Research Support, Non-U.S. Gov’t 2016/02/18 ACS Synth Biol. 2016 May 20;5(5):395–404. doi: 10.1021/acssynbio.5b00266. Epub 2016 Mar 1.

Faulkner NE, Dujardin DL, Tai CY, Vaughan KT, O’Connell CB, Wang Y, Vallee RB. A role for the lissencephaly gene LIS1 in mitosis and cytoplasmic dynein function. Nat Cell Biol. 2000; 2(11):784–91. https://www.ncbi.nlm.nih.gov/pubmed/11056532, doi: 10.1038/35041020, faulkner, E E Dujardin, L L Tai, Y Y Vaughan, K T O’Connell, B B Wang, Y Vallee, R B eng GM47434/GM/NIGMS NIH HHS/HD61982/HD/NICHD NIH HHS/Research Support, Non-U.S. Gov’t Research Support, U.S. Gov’t, P.H.S. England 2000/11/01 Nat Cell Biol. 2000 Nov;2(11):784–91. doi: 10.1038/35041020.

Gerlach M, Kraft T, Brenner B, Petersen B, Niemann H, Montag J. Efficient Knock-in of a Point Mutation in Porcine Fibroblasts Using the CRISPR/Cas9-GMNN Fusion Gene. Genes (Basel). 2018; 9(6). https://www.ncbi.nlm.nih.gov/pubmed/29899280, doi: 10.3390/genes9060296, gerlach, Max Kraft, Theresia Brenner, Bernhard Petersen, Bjorn Niemann, Heiner Montag, Judith eng Switzerland 2018/06/15 Genes (Basel). 2018 Jun 13;9(6):296. doi: 10.3390/genes9060296.

Ghetti S, Burigotto M, Mattivi A, Magnani G, Casini A, Bianchi A, Cereseto A, Fava LL. CRISPR/Cas9 ribonucleoprotein-mediated knockin generation in hTERT-RPE1 cells. STAR Protoc. 2021; 2(2):100407. https://www.ncbi.nlm.nih.gov/pubmed/33855309, doi: 10.1016/j.xpro.2021.100407, ghetti, Sabrina Burigotto, Matteo Mattivi, Alessia Magnani, Giovanni Casini, Antonio Bianchi, Andrea Cereseto, Anna Fava, Luca L eng Research Support, Non-U.S. Gov’t 2021/04/16 STAR Protoc. 2021 Mar 24;2(2):100407. doi: 10.1016/j.xpro.2021.100407. eCollection 2021 Jun 18.

Han Z, Huang C, Luo T, Mirkin CA. A general genome editing strategy using CRISPR lipid nanoparticle spherical nucleic acids. Proc Natl Acad Sci U S A. 2025; 122(36):e2426094122. https://www.ncbi.nlm.nih.gov/pubmed/40906807, doi: 10.1073/pnas.2426094122, han, Zhenyu Huang, Chi Luo, Taokun Mirkin, Chad A eng FA9550-22-1-0300/DOD | AF | AMC | AFRL | Air Force Office of Scientific Research (AFOSR)/DMR-2428112/National Science Foundation (NSF)/DMR-2308691/National Science Foundation (NSF)/NSF ECCS-2025633/National Science Foundation (NSF)/2025/09/04 Proc Natl Acad Sci U S A. 2025 Sep 9;122(36):e2426094122. doi: 10.1073/pnas.2426094122. Epub 2025 Sep 4.

Harada A, Takei Y, Kanai Y, Tanaka Y, Nonaka S, Hirokawa N. Golgi vesiculation and lysosome dispersion in cells lacking cytoplasmic dynein. J Cell Biol. 1998; 141(1):51–9. https://www.ncbi.nlm.nih.gov/pubmed/9531547, doi: 10.1083/jcb.141.1.51, harada, A Takei, Y Kanai, Y Tanaka, Y Nonaka, S Hirokawa, N eng N01-HD-2-3144/HD/NICHD NIH HHS/Research Support, Non-U.S. Gov’t Research Support, U.S. Gov’t, P.H.S. 1998/05/16 J Cell Biol. 1998 Apr 6;141(1):51–9. doi: 10.1083/jcb.141.1.51.

Howell BJ, McEwen BF, Canman JC, Hoffman DB, Farrar EM, Rieder CL, Salmon ED. Cytoplasmic dynein/dynactin drives kinetochore protein transport to the spindle poles and has a role in mitotic spindle checkpoint inactivation. J Cell Biol. 2001; 155(7):1159–72. https://www.ncbi.nlm.nih.gov/pubmed/11756470, doi: 10.1083/jcb.200105093, howell, J J McEwen, F F Canman, C C Hoffman, B B Farrar, M M Rieder, L L Salmon, E D eng GM-24364/GM/NIGMS NIH HHS/GMS-R01-40198/GM/NIGMS NIH HHS/R37 GM040198/GM/NIGMS NIH HHS/R37 GM024364/GM/NIGMS NIH HHS/R01 GM024364/GM/NIGMS NIH HHS/Research Support, U.S. Gov’t, Non-P.H.S. Research Support, U.S. Gov’t, P.H.S. 2002/01/05 J Cell Biol. 2001 Dec 24;155(7):1159–72. doi: 10.1083/jcb.200105093. Epub 2001 Dec 24.

Iyer S, Mir A, Vega-Badillo J, Roscoe BP, Ibraheim R, Zhu LJ, Lee J, Liu P, Luk K, Mintzer E, Guo D, Soares de Brito J, Emerson J C P, Zamore PD, Sontheimer EJ, Wolfe SA. Efficient Homology-Directed Repair with Circular Single-Stranded DNA Donors. CRISPR J. 2022; 5(5):685–701. https://www.ncbi.nlm.nih.gov/pubmed/36070530, doi: 10.1089/crispr.2022.0058, iyer, Sukanya Mir, Aamir Vega-Badillo, Joel Roscoe, Benjamin P Ibraheim, Raed Zhu, Lihua Julie Lee, Jooyoung Liu, Pengpeng Luk, Kevin Mintzer, Esther Guo, Dongsheng Soares de Brito, Josias Emerson, Charles P Jr Zamore, Phillip D Sontheimer, Erik J Wolfe, Scot A eng Research Support, N.I.H., Extramural 2022/09/08 CRISPR J. 2022 Oct;5(5):685–701. doi: 10.1089/crispr.2022.0058. Epub 2022 Sep 7.

Jiang XR, Jimenez G, Chang E, Frolkis M, Kusler B, Sage M, Beeche M, Bodnar AG, Wahl GM, Tlsty TD, Chiu CP. Telomerase expression in human somatic cells does not induce changes associated with a transformed phenotype. Nat Genet. 1999; 21(1):111–4. https://www.ncbi.nlm.nih.gov/pubmed/9916802, doi: 10.1038/5056, jiang, R R Jimenez, G Chang, E Frolkis, M Kusler, B Sage, M Beeche, M Bodnar, G G Wahl, M M Tlsty, D D Chiu, C P eng CA 42765/CA/NCI NIH HHS/CA 51912/CA/NCI NIH HHS/CA 61449/CA/NCI NIH HHS/etc. Research Support, Non-U.S. Gov’t Research Support, U.S. Gov’t, P.H.S. 1999/01/23 Nat Genet. 1999 Jan;21(1):111–4. doi: 10.1038/5056.

Jin YY, Zhang P, Liu DP. Optimizing homology-directed repair for gene editing: the potential of single-stranded DNA donors. Trends Genet. 2025; 41(9):788–803. https://www.ncbi.nlm.nih.gov/pubmed/40436684, doi: 10.1016/j.tig.2025.04.014, jin, Ying-Ying Zhang, Peng Liu, De-Pei eng Review England 2025/05/29 Trends Genet. 2025 Sep;41(9):788–803. doi: 10.1016/j.tig.2025.04.014. Epub 2025 May 27.

Jinek M, Chylinski K, Fonfara I, Hauer M, Doudna JA, Charpentier E. A programmable dual-RNA-guided DNA endonuclease in adaptive bacterial immunity. Science. 2012; 337(6096):816–21. https://www.ncbi.nlm.nih.gov/pubmed/22745249, doi: 10.1126/science.1225829, jinek, Martin Chylinski, Krzysztof Fonfara, Ines Hauer, Michael Doudna, Jennifer A Charpentier, Emmanuelle eng HHMI/Howard Hughes Medical Institute/Research Support, Non-U.S. Gov’t 2012/06/30 Science. 2012 Aug 17;337(6096):816–21. doi: 10.1126/science.1225829. Epub 2012 Jun 28.

Kadonaga JT. Perspectives on the RNA polymerase II core promoter. Wiley Interdiscip Rev Dev Biol. 2012; 1(1):40–51. https://www.ncbi.nlm.nih.gov/pubmed/23801666, doi: 10.1002/wdev.21, kadonaga, James T eng R01 GM041249/GM/NIGMS NIH HHS/GM041249/GM/NIGMS NIH HHS/Research Support, N.I.H., Extramural Research Support, Non-U.S. Gov’t Review 2012/01/01 Wiley Interdiscip Rev Dev Biol. 2012 Jan-Feb;1(1):40–51. doi: 10.1002/wdev.21. Epub 2011 Dec 6.

Kanie T, Abbott KL, Mooney NA, Plowey ED, Demeter J, Jackson PK. The CEP19-RABL2 GTPase Complex Binds IFT-B to Initiate Intraflagellar Transport at the Ciliary Base. Dev Cell. 2017; 42(1):22–36 e12. https://www.ncbi.nlm.nih.gov/pubmed/28625565, doi: 10.1016/j.devcel.2017.05.016, kanie, Tomoharu Abbott, Keene Louis Mooney, Nancie Ann Plowey, Edward Douglas Demeter, Janos Jackson, Peter Kent eng K08 NS085324/NS/NINDS NIH HHS/R01 GM114276/GM/NIGMS NIH HHS/R01 GM121565/GM/NIGMS NIH HHS/Research Support, N.I.H., Extramural Research Support, Non-U.S. Gov’t 2017/06/20 Dev Cell. 2017 Jul 10;42(1):22–36.e12. doi: 10.1016/j.devcel.2017.05.016. Epub 2017 Jun 15.

Kanie T, Ng R, Abbott KL, Tanvir NM, Lorentzen E, Pongs O, Jackson PK. Myristoylated Neuronal Calcium Sensor-1 captures the preciliary vesicle at distal appendages. Elife. 2025; 14. https://www.ncbi.nlm.nih.gov/pubmed/39882855, doi: 10.7554/eLife.85998, kanie, Tomoharu Ng, Roy Abbott, Keene L Tanvir, Niaj Mohammad Lorentzen, Esben Pongs, Olaf Jackson, Peter K eng P20 GM103639/GM/NIGMS NIH HHS/P20GM103447/GM/NIGMS NIH HHS/P30 CA225520/CA/NCI NIH HHS/R01 GM121565/GM/NIGMS NIH HHS/R01 GM114276/GM/NIGMS NIH HHS/R01GM114276/GM/NIGMS NIH HHS/R35 GM151013/GM/NIGMS NIH HHS/R01GM121565/GM/NIGMS NIH HHS/S10 OD028536/OD/NIH HHS/P20 GM103447/GM/NIGMS NIH HHS/1R35GM151013/GM/NIGMS NIH HHS/England 2025/01/30 Elife. 2025 Jan 30;14:e85998. doi: 10.7554/eLife.85998.

Kim DW, Uetsuki T, Kaziro Y, Yamaguchi N, Sugano S. Use of the human elongation factor 1 alpha promoter as a versatile and efficient expression system. Gene. 1990; 91(2):217–23. https://www.ncbi.nlm.nih.gov/pubmed/2210382, doi: 10.1016/0378-1119(90)90091-5, kim, W W Uetsuki, T Kaziro, Y Yamaguchi, N Sugano, S eng Research Support, Non-U.S. Gov’t Netherlands 1990/07/16 Gene. 1990 Jul 16;91(2):217–23. doi: 10.1016/0378-1119(90)90091-5.

Koduri V, McBrayer SK, Liberzon E, Wang AC, Briggs KJ, Cho H, Kaelin J W G. Peptidic degron for IMiD-induced degradation of heterologous proteins. Proc Natl Acad Sci U S A. 2019; 116(7):2539–2544. https://www.ncbi.nlm.nih.gov/pubmed/30683719, doi: 10.1073/pnas.1818109116, koduri, Vidyasagar McBrayer, Samuel K Liberzon, Ella Wang, Adam C Briggs, Kimberly J Cho, Hyejin Kaelin, William G Jr eng R01 CA068490/CA/NCI NIH HHS/P50 CA101942/CA/NCI NIH HHS/HHMI/Howard Hughes Medical Institute/R35 CA210068/CA/NCI NIH HHS/T32 CA009172/CA/NCI NIH HHS/P30 CA006516/CA/NCI NIH HHS/P50 CA165962/CA/NCI NIH HHS/Research Support, N.I.H., Extramural Research Support, Non-U.S. Gov’t 2019/01/27 Proc Natl Acad Sci U S A. 2019 Feb 12;116(7):2539–2544. doi: 10.1073/pnas.1818109116. Epub 2019 Jan 25.

Kurashina M, Mizumoto K. Targeting endogenous proteins for spatial and temporal knockdown using auxin-inducible degron in Caenorhabditis elegans. STAR Protoc. 2023; 4(1):102028. https://www.ncbi.nlm.nih.gov/pubmed/36640369, doi: 10.1016/j.xpro.2022.102028, kurashina, Mizuki Mizumoto, Kota eng P40 OD010440/OD/NIH HHS/PJT-148667/CIHR/Canada Research Support, N.I.H., Extramural Research Support, Non-U.S. Gov’t 2023/01/15 STAR Protoc. 2023 Mar 17;4(1):102028. doi: 10.1016/j.xpro.2022.102028. Epub 2023 Jan 13.

Larsson AJM, Johnsson P, Hagemann-Jensen M, Hartmanis L, Faridani OR, Reinius B, Segerstolpe A, Rivera CM, Ren B, Sandberg R. Genomic encoding of transcriptional burst kinetics. Nature. 2019; 565(7738):251–254. https://www.ncbi.nlm.nih.gov/pubmed/30602787, doi: 10.1038/s41586-018-0836-1, larsson, Anton M M Johnsson, Per Hagemann-Jensen, Michael Hartmanis, Leonard Faridani, Omid R Reinius, Bjorn Segerstolpe, Asa Rivera, Chloe M Ren, Bing Sandberg, Rickard eng 648842/ERC_/European Research Council/International Research Support, Non-U.S. Gov’t England 2019/01/04 Nature. 2019 Jan;565(7738):251–254. doi: 10.1038/s41586-018-0836-1. Epub 2019 Jan 2.

Letort G, Duclert A, Le Clerre D, Chion-Sotinel I, Salvatori R, Dessez E, Sevin M, Rotondi M, Ducani C, Duchateau P, Valton J. Circular single stranded DNA potentiates non-viral gene insertion in hematopoietic stem and progenitor cells. Nat Commun. 2025; 16(1):10125. https://www.ncbi.nlm.nih.gov/pubmed/41257973, doi: 10.1038/s41467-025-66318-2, letort, Gil Duclert, Aymeric Le Clerre, Diane Chion-Sotinel, Isabelle Salvatori, Roger Dessez, Emilie Sevin, Margaux Rotondi, Marco Ducani, Cosimo Duchateau, Philippe Valton, Julien eng England 2025/11/19 Nat Commun. 2025 Nov 19;16(1):10125. doi: 10.1038/s41467-025-66318-2.

Li J, Liang Q, Zhou H, Zhou M, Huang H. Profiling the impact of the promoters on CRISPR-Cas12a system in human cells. Cell Mol Biol Lett. 2023; 28(1):41. https://www.ncbi.nlm.nih.gov/pubmed/37198545, doi: 10.1186/s11658-023-00454-9, li, Jinhe Liang, Qinchun Zhou, HuaPing Zhou, Ming Huang, Hongxin eng Letter England 2023/05/18 Cell Mol Biol Lett. 2023 May 17;28(1):41. doi: 10.1186/s11658-023-00454-9.

Li K, Wang G, Andersen T, Zhou P, Pu WT. Optimization of genome engineering approaches with the CRISPR/Cas9 system. PLoS One. 2014; 9(8):e105779. https://www.ncbi.nlm.nih.gov/pubmed/25166277, doi: 10.1371/journal.pone.0105779, li, Kai Wang, Gang Andersen, Troels Zhou, Pingzhu Pu, William T eng U01 HL098188/HL/NHLBI NIH HHS/U01HL098188/HL/NHLBI NIH HHS/T32 HL007572/HL/NHLBI NIH HHS/U01HL098166/HL/NHLBI NIH HHS/U01 HL098166/HL/NHLBI NIH HHS/Research Support, N.I.H., Extramural 2014/08/29 PLoS One. 2014 Aug 28;9(8):e105779. doi: 10.1371/journal.pone.0105779. eCollection 2014.

Li Z, Ran H, Tang X, Liu X, Wan X, Xiong P, Gan Z, Liu X, Wang L, Yuan J. Proteomics Reveals AP-2 Complex Depletion Suppressing Listeria monocytogenes Intracellular Replication. Proteomics. 2025; 25(19):26–38. https://www.ncbi.nlm.nih.gov/pubmed/40847839, doi: 10.1002/pmic.70034, li, Zhangfu Ran, Haiying Tang, Xiangyu Liu, Xiao Wan, Xiaoyuan Xiong, Pei Gan, Zhe Liu, Xu Wang, Liting Yuan, Jiangbei eng 82402038/National Natural Science Foundation of China/2024A1515013277/Guangdong Basic and Applied Basic Research Foundation/JCYJ20230807095808017/Shenzhen Science and Technology Program/Germany 2025/08/23 Proteomics. 2025 Oct;25(19):26–38. doi: 10.1002/pmic.70034. Epub 2025 Aug 23.

Lin S, Staahl BT, Alla RK, Doudna JA. Enhanced homology-directed human genome engineering by controlled timing of CRISPR/Cas9 delivery. Elife. 2014; 3:e04766. https://www.ncbi.nlm.nih.gov/pubmed/25497837, doi: 10.7554/eLife.04766, lin, Steven Staahl, Brett T Alla, Ravi K Doudna, Jennifer A eng Research Support, Non-U.S. Gov’t England 2014/12/17 Elife. 2014 Dec 15;3:e04766. doi: 10.7554/eLife.04766.

Liu Z, Chen O, Wall JBJ, Zheng M, Zhou Y, Wang L, Vaseghi HR, Qian L, Liu J. Systematic comparison of 2A peptides for cloning multi-genes in a polycistronic vector. Sci Rep. 2017; 7(1):2193. https://www.ncbi.nlm.nih.gov/pubmed/28526819, doi: 10.1038/s41598-017-02460-2, liu, Ziqing Chen, Olivia Wall, J Blake Joseph Zheng, Michael Zhou, Yang Wang, Li Vaseghi, Haley Ruth Qian, Li Liu, Jiandong eng R00 HL109079/HL/NHLBI NIH HHS/R01 HL128331/HL/NHLBI NIH HHS/Research Support, N.I.H., Extramural Research Support, Non-U.S. Gov’t England 2017/05/21 Sci Rep. 2017 May 19;7(1):2193. doi: 10.1038/s41598-017-02460-2.

Lujan P, Garcia-Cabau C, Wakana Y, Vera Lillo J, Rodilla-Ramirez C, Sugiura H, Malhotra V, Salvatella X, Garcia-Parajo MF, Campelo F. Sorting of secretory proteins at the trans-Golgi network by human TGN46. Elife. 2024; 12. https://www.ncbi.nlm.nih.gov/pubmed/38466628, doi: 10.7554/eLife.91708, lujan, Pablo Garcia-Cabau, Carla Wakana, Yuichi Vera Lillo, Javier Rodilla-Ramirez, Carmen Sugiura, Hideaki Malhotra, Vivek Salvatella, Xavier Garcia-Parajo, Maria F Campelo, Felix eng CONCERT (GA 648201)/ERC_/European Research Council/International NANO-MEMEC (GA 788546)/ERC_/European Research Council/International England 2024/03/11 Elife. 2024 Mar 11;12:RP91708. doi: 10.7554/eLife.91708.

Maneiro MA, Forte N, Shchepinova MM, Kounde CS, Chudasama V, Baker JR, Tate EW. Antibody-PROTAC Conjugates Enable HER2-Dependent Targeted Protein Degradation of BRD4. ACS Chem Biol. 2020; 15(6):1306–1312. https://www.ncbi.nlm.nih.gov/pubmed/32338867, doi: 10.1021/acschembio.0c00285, maneiro, Mari A Forte, Nafsika Shchepinova, Maria M Kounde, Cyrille S Chudasama, Vijay Baker, James Richard Tate, Edward W eng Research Support, Non-U.S. Gov’t 2020/04/28 ACS Chem Biol. 2020 Jun 19;15(6):1306–1312. doi: 10.1021/acschembio.0c00285. Epub 2020 Apr 30.

Maruyama T, Dougan SK, Truttmann MC, Bilate AM, Ingram JR, Ploegh HL. Increasing the efficiency of precise genome editing with CRISPR-Cas9 by inhibition of nonhomologous end joining. Nat Biotechnol. 2015; 33(5):538–42. https://www.ncbi.nlm.nih.gov/pubmed/25798939, doi: 10.1038/nbt.3190, maruyama, Takeshi Dougan, Stephanie K Truttmann, Matthias C Bilate, Angelina M Ingram, Jessica R Ploegh, Hidde L eng R01 AI087879/AI/NIAID NIH HHS/R01 AI087879-01/AI/NIAID NIH HHS/Research Support, N.I.H., Extramural Research Support, Non-U.S. Gov’t 2015/03/24 Nat Biotechnol. 2015 May;33(5):538–42. doi: 10.1038/nbt.3190. Epub 2015 Mar 23.

Miyamoto T, Hosoba K, Ochiai H, Royba E, Izumi H, Sakuma T, Yamamoto T, Dynlacht BD, Matsuura S. The Microtubule-Depolymerizing Activity of a Mitotic Kinesin Protein KIF2A Drives Primary Cilia Disassembly Coupled with Cell Proliferation. Cell Rep. 2015; 10(5):664–673. https://www.ncbi.nlm.nih.gov/pubmed/25660017, doi: 10.1016/j.celrep.2015.01.003, miyamoto, Tatsuo Hosoba, Kosuke Ochiai, Hiroshi Royba, Ekaterina Izumi, Hideki Sakuma, Tetsushi Yamamoto, Takashi Dynlacht, Brian David Matsuura, Shinya eng R01 HD069647/HD/NICHD NIH HHS/2015/02/11 Cell Rep. 2015 Feb 10;10(5):664–673. doi: 10.1016/j.celrep.2015.01.003. Epub 2015 Feb 5.

Miyaoka Y, Berman JR, Cooper SB, Mayerl SJ, Chan AH, Zhang B, Karlin-Neumann GA, Conklin BR. Systematic quantification of HDR and NHEJ reveals effects of locus, nuclease, and cell type on genome-editing. Sci Rep. 2016; 6:23549. https://www.ncbi.nlm.nih.gov/pubmed/27030102, doi: 10.1038/srep23549, miyaoka, Yuichiro Berman, Jennifer R Cooper, Samantha B Mayerl, Steven J Chan, Amanda H Zhang, Bin Karlin-Neumann, George A Conklin, Bruce R eng P01-HL089707/HL/NHLBI NIH HHS/U01 HL099997/HL/NHLBI NIH HHS/R01 HL060664/HL/NHLBI NIH HHS/U01-HL100406/HL/NHLBI NIH HHS/P01 HL089707/HL/NHLBI NIH HHS/U01 HL098179/HL/NHLBI NIH HHS/R01-HL060664/HL/NHLBI NIH HHS/U01-GM09614/GM/NIGMS NIH HHS/U01-HL098179/HL/NHLBI NIH HHS/U01-HL099997/HL/NHLBI NIH HHS/U01 HL100406/HL/NHLBI NIH HHS/R01 HL108677/HL/NHLBI NIH HHS/R01-HL108677/HL/NHLBI NIH HHS/R01 HL130533/HL/NHLBI NIH HHS/Research Support, N.I.H., Extramural Research Support, Non-U.S. Gov’t England 2016/04/01 Sci Rep. 2016 Mar 31;6:23549. doi: 10.1038/srep23549.

Motley A, Bright NA, Seaman MN, Robinson MS. Clathrin-mediated endocytosis in AP-2-depleted cells. J Cell Biol. 2003; 162(5):909–18. https://www.ncbi.nlm.nih.gov/pubmed/12952941, doi: 10.1083/jcb.200305145, motley, Alison Bright, Nicholas A Seaman, MatthewJ J Robinson, Margaret S eng Wellcome Trust/United Kingdom Research Support, Non-U.S. Gov’t 2003/09/04 J Cell Biol. 2003 Sep 1;162(5):909–18. doi: 10.1083/jcb.200305145.

Nabet B, Ferguson FM, Seong BKA, Kuljanin M, Leggett AL, Mohardt ML, Robichaud A, Conway AS, Buckley DL, Mancias JD, Bradner JE, Stegmaier K, Gray NS. Rapid and direct control of target protein levels with VHL-recruiting dTAG molecules. Nat Commun. 2020; 11(1):4687. https://www.ncbi.nlm.nih.gov/pubmed/32948771, doi: 10.1038/s41467-020-18377-w, nabet, Behnam Ferguson, Fleur M Seong, Bo Kyung A Kuljanin, Miljan Leggett, Alan L Mohardt, Mikaela L Robichaud, Amanda Conway, Amy S Buckley, Dennis L Mancias, Joseph D Bradner, James E Stegmaier, Kimberly Gray, Nathanael S eng R01 CA204915/CA/NCI NIH HHS/U54 CA231637/CA/NCI NIH HHS/Research Support, N.I.H., Extramural Research Support, Non-U.S. Gov’t Research Support, U.S. Gov’t, Non-P.H.S. England 2020/09/20 Nat Commun. 2020 Sep 18;11(1):4687. doi: 10.1038/s41467-020-18377-w.

Nabet B, Roberts JM, Buckley DL, Paulk J, Dastjerdi S, Yang A, Leggett AL, Erb MA, Lawlor MA, Souza A, Scott TG, Vittori S, Perry JA, Qi J, Winter GE, Wong KK, Gray NS, Bradner JE. The dTAG system for immediate and target-specific protein degradation. Nat Chem Biol. 2018; 14(5):431–441. https://www.ncbi.nlm.nih.gov/pubmed/29581585, doi: 10.1038/s41589-018-0021-8, nabet, Behnam Roberts, Justin M Buckley, Dennis L Paulk, Joshiawa Dastjerdi, Shiva Yang, Annan Leggett, Alan L Erb, Michael A Lawlor, Matthew A Souza, Amanda Scott, Thomas G Vittori, Sarah Perry, Jennifer A Qi, Jun Winter, Georg E Wong, Kwok-Kin Gray, Nathanael S Bradner, James E eng P01 CA066996/CA/NCI NIH HHS/R01 CA140594/CA/NCI NIH HHS/R01 CA166480/CA/NCI NIH HHS/Research Support, Non-U.S. Gov’t 2018/03/28 Nat Chem Biol. 2018 May;14(5):431–441. doi: 10.1038/s41589-018-0021-8. Epub 2018 Mar 26.

Nishimura K, Fukagawa T, Takisawa H, Kakimoto T, Kanemaki M. An auxin-based degron system for the rapid depletion of proteins in nonplant cells. Nat Methods. 2009; 6(12):917–22. https://www.ncbi.nlm.nih.gov/pubmed/19915560, doi: 10.1038/nmeth.1401, nishimura, Kohei Fukagawa, Tatsuo Takisawa, Haruhiko Kakimoto, Tatsuo Kanemaki, Masato eng Research Support, Non-U.S. Gov’t 2009/11/17 Nat Methods. 2009 Dec;6(12):917–22. doi: 10.1038/nmeth.1401. Epub 2009 Nov 15.

Palmer KJ, Hughes H, Stephens DJ. Specificity of cytoplasmic dynein subunits in discrete membrane-trafficking steps. Mol Biol Cell. 2009; 20(12):2885–99. https://www.ncbi.nlm.nih.gov/pubmed/19386764, doi: 10.1091/mbc.e08-12-1160, palmer, Krysten J Hughes, Helen Stephens, David J eng G117/554/MRC_/Medical Research Council/United Kingdom G117/554(71630)/MRC_/Medical Research Council/United Kingdom G117/553/MRC_/Medical Research Council/United Kingdom BB_/Biotechnology and Biological Sciences Research Council/United Kingdom Research Support, Non-U.S. Gov’t 2009/04/24 Mol Biol Cell. 2009 Jun;20(12):2885–99. doi: 10.1091/mbc.e08-12-1160. Epub 2009 Apr 22.

Philip R, Sharma A, Matellan L, Erpf AC, Hsu WH, Tkach JM, Wyatt HDM, Pelletier L. qTAG: an adaptable plasmid scaffold for CRISPR-based endogenous tagging. EMBO J. 2025; 44(3):947–974. https://www.ncbi.nlm.nih.gov/pubmed/39668248, doi: 10.1038/s44318-024-00337-5, philip, Reuben Sharma, Amit Matellan, Laura Erpf, Anna C Hsu, Wen-Hsin Tkach, Johnny M Wyatt, HaleyM M Pelletier, Laurence eng 187836/Canadian Government | CIHR | Institute of Cancer Research (IC)/181763/Canadian Government | CIHR | Institute of Cancer Research (IC)/156297/Canadian Government | Canadian Institutes of Health Research (CIHR)/167279/Canadian Government | Canadian Institutes of Health Research (CIHR)/N/A/Krembil Foundation/England 2024/12/13 EMBO J. 2025 Feb;44(3):947–974. doi: 10.1038/s44318-024-00337-5. Epub 2024 Dec 12.

Pillow TH, Adhikari P, Blake RA, Chen J, Del Rosario G, Deshmukh G, Figueroa I, Gascoigne KE, Kamath AV, Kaufman S, Kleinheinz T, Kozak KR, Latifi B, Leipold DD, Sing Li C, Li R, Mulvihill MM, O’Donohue A, Rowntree RK, Sadowsky JD, et al. Antibody Conjugation of a Chimeric BET Degrader Enables in vivo Activity. ChemMedChem. 2020; 15(1):17–25. https://www.ncbi.nlm.nih.gov/pubmed/31674143, doi: 10.1002/cmdc.201900497, pillow, Thomas H Adhikari, Pragya Blake, Robert A Chen, Jinhua Del Rosario, Geoffrey Deshmukh, Gauri Figueroa, Isabel Gascoigne, Karen E Kamath, Amrita V Kaufman, Susan Kleinheinz, Tracy Kozak, Katherine R Latifi, Brandon Leipold, Douglas D Sing Li, Chun Li, Ruina Mulvihill, Melinda M O’Donohue, Aimee Rowntree, Rebecca K Sadowsky, Jack D Wai, John Wang, Xinxin Wu, Cong Xu, Zijin Yao, Hui Yu, Shang-Fan Zhang, Donglu Zang, Richard Zhang, Hongyan Zhou, Hao Zhu, Xiaoyu Dragovich, Peter S eng Germany 2019/11/02 ChemMedChem. 2020 Jan 7;15(1):17–25. doi: 10.1002/cmdc.201900497. Epub 2019 Nov 14.

Pinder J, Salsman J, Dellaire G. Nuclear domain ‘knock-in’ screen for the evaluation and identification of small molecule enhancers of CRISPR-based genome editing. Nucleic Acids Res. 2015; 43(19):9379–92. https://www.ncbi.nlm.nih.gov/pubmed/26429972, doi: 10.1093/nar/gkv993, pinder, Jordan Salsman, Jayme Dellaire, Graham eng MOP-84260/Canadian Institutes of Health Research/Canada Research Support, Non-U.S. Gov’t England 2015/10/03 Nucleic Acids Res. 2015 Oct 30;43(19):9379–92. doi: 10.1093/nar/gkv993. Epub 2015 Oct 1.

Raaijmakers JA, Tanenbaum ME, Medema RH. Systematic dissection of dynein regulators in mitosis. J Cell Biol. 2013; 201(2):201–15. https://www.ncbi.nlm.nih.gov/pubmed/23589491, doi: 10.1083/jcb.201208098, raaijmakers, Jonne A Tanenbaum, Marvin E Medema, Rene H eng Research Support, Non-U.S. Gov’t 2013/04/17 J Cell Biol. 2013 Apr 15;201(2):201–15. doi: 10.1083/jcb.201208098.

Renaud JB, Boix C, Charpentier M, De Cian A, Cochennec J, Duvernois-Berthet E, Perrouault L, Tesson L, Edouard J, Thinard R, Cherifi Y, Menoret S, Fontaniere S, de Croze N, Fraichard A, Sohm F, Anegon I, Concordet JP, Giovannangeli C. Improved Genome Editing Efficiency and Flexibility Using Modified Oligonucleotides with TALEN and CRISPR-Cas9 Nucleases. Cell Rep. 2016; 14(9):2263–2272. https://www.ncbi.nlm.nih.gov/pubmed/26923600, doi: 10.1016/j.celrep.2016.02.018, renaud, Jean-Baptiste Boix, Charlotte Charpentier, Marine De Cian, Anne Cochennec, Julien Duvernois-Berthet, Evelyne Perrouault, Loic Tesson, Laurent Edouard, Joanne Thinard, Reynald Cherifi, Yacine Menoret, Severine Fontaniere, Sandra de Croze, Noemie Fraichard, Alexandre Sohm, Frederic Anegon, Ignacio Concordet, Jean-Paul Giovannangeli, Carine eng Research Support, Non-U.S. Gov’t 2016/03/01 Cell Rep. 2016 Mar 8;14(9):2263–2272. doi: 10.1016/j.celrep.2016.02.018. Epub 2016 Feb 25.

Riesenberg S, Maricic T. Targeting repair pathways with small molecules increases precise genome editing in pluripotent stem cells. Nat Commun. 2018; 9(1):2164. https://www.ncbi.nlm.nih.gov/pubmed/29867139, doi: 10.1038/s41467-018-04609-7, riesenberg, Stephan Maricic, Tomislav eng Research Support, Non-U.S. Gov’t England 2018/06/06 Nat Commun. 2018 Jun 4;9(1):2164. doi: 10.1038/s41467-018-04609-7.

Roth TL, Puig-Saus C, Yu R, Shifrut E, Carnevale J, Li PJ, Hiatt J, Saco J, Krystofinski P, Li H, Tobin V, Nguyen DN, Lee MR, Putnam AL, Ferris AL, Chen JW, Schickel JN, Pellerin L, Carmody D, Alkorta-Aranburu G, et al. Reprogramming human T cell function and specificity with non-viral genome targeting. Nature. 2018; 559(7714):405–409. https://www.ncbi.nlm.nih.gov/pubmed/29995861, doi: 10.1038/s41586-018-0326-5, roth, Theodore L Puig-Saus, Cristina Yu, Ruby Shifrut, Eric Carnevale, Julia Li, P Jonathan Hiatt, Joseph Saco, Justin Krystofinski, Paige Li, Han Tobin, Victoria Nguyen, David N Lee, Michael R Putnam, Amy L Ferris, Andrea L Chen, Jeff W Schickel, Jean-Nicolas Pellerin, Laurence Carmody, David Alkorta-Aranburu, Gorka Del Gaudio, Daniela Matsumoto, Hiroyuki Morell, Montse Mao, Ying Cho, Min Quadros, Rolen M Gurumurthy, Channabasavaiah B Smith, Baz Haugwitz, Michael Hughes, Stephen H Weissman, Jonathan S Schumann, Kathrin Esensten, Jonathan H May, Andrew P Ashworth, Alan Kupfer, Gary M Greeley, Siri Atma W Bacchetta, Rosa Meffre, Eric Roncarolo, Maria Grazia Romberg, Neil Herold, Kevan C Ribas, Antoni Leonetti, Manuel D Marson, Alexander eng T32 GM007618/GM/NIGMS NIH HHS/L40 AI140341/AI/NIAID NIH HHS/P30 CA082103/CA/NCI NIH HHS/R35 CA197633/CA/NCI NIH HHS/S10 OD021822/OD/NIH HHS/T32 AI007641/AI/NIAID NIH HHS/P50 GM082250/GM/NIGMS NIH HHS/T32 CA108462/CA/NCI NIH HHS/K23 DK094866/DK/NIDDK NIH HHS/K23 AI115001/AI/NIAID NIH HHS/P30 DK020595/DK/NIDDK NIH HHS/P30 DK063720/DK/NIDDK NIH HHS/DP3 DK111914/DK/NIDDK NIH HHS/T32 AI007334/AI/NIAID NIH HHS/T32 DK007418/DK/NIDDK NIH HHS/Research Support, N.I.H., Extramural Research Support, Non-U.S. Gov’t England 2018/07/12 Nature. 2018 Jul;559(7714):405–409. doi: 10.1038/s41586-018-0326-5. Epub 2018 Jul 11.

Savic N, Ringnalda FC, Lindsay H, Berk C, Bargsten K, Li Y, Neri D, Robinson MD, Ciaudo C, Hall J, Jinek M, Schwank G. Covalent linkage of the DNA repair template to the CRISPR-Cas9 nuclease enhances homology-directed repair. Elife. 2018; 7. https://www.ncbi.nlm.nih.gov/pubmed/29809142, doi: 10.7554/eLife.33761, savic, Natasa Ringnalda, Femke Cas Lindsay, Helen Berk, Christian Bargsten, Katja Li, Yizhou Neri, Dario Robinson, Mark D Ciaudo, Constance Hall, Jonathan Jinek, Martin Schwank, Gerald eng PMPDP3_171388/Swiss National Science Foundation/Switzerland 31003A_160230/Swiss National Science Foundation/Switzerland 31003A_149393/Swiss National Science Foundation/Switzerland Research Support, Non-U.S. Gov’t England 2018/05/29 Elife. 2018 May 29;7:e33761. doi: 10.7554/eLife.33761.

Schumann K, Lin S, Boyer E, Simeonov DR, Subramaniam M, Gate RE, Haliburton GE, Ye CJ, Bluestone JA, Doudna JA, Marson A. Generation of knock-in primary human T cells using Cas9 ribonucleoproteins. Proc Natl Acad Sci U S A. 2015; 112(33):10437–42. https://www.ncbi.nlm.nih.gov/pubmed/26216948, doi: 10.1073/pnas.1512503112, schumann, Kathrin Lin, Steven Boyer, Eric Simeonov, Dimitre R Subramaniam, Meena Gate, Rachel E Haliburton, Genevieve E Ye, Chun J Bluestone, Jeffrey A Doudna, Jennifer A Marson, Alexander eng P30 DK063720/DK/NIDDK NIH HHS/DRG-(2176-13)/Howard Hughes Medical Institute/T32 GM008284/GM/NIGMS NIH HHS/T32 GM067547/GM/NIGMS NIH HHS/T32 DK741834/DK/NIDDK NIH HHS/P50GM082250/GM/NIGMS NIH HHS/T32 EB009383/EB/NIBIB NIH HHS/P50 GM082250/GM/NIGMS NIH HHS/T32 GM008568/GM/NIGMS NIH HHS/Research Support, N.I.H., Extramural Research Support, Non-U.S. Gov’t 2015/07/29 Proc Natl Acad Sci U S A. 2015 Aug 18;112(33):10437–42. doi: 10.1073/pnas.1512503112. Epub 2015 Jul 27.

Scully R, Panday A, Elango R, Willis NA. DNA double-strand break repair-pathway choice in somatic mammalian cells. Nat Rev Mol Cell Biol. 2019; 20(11):698–714. https://www.ncbi.nlm.nih.gov/pubmed/31263220, doi: 10.1038/s41580-019-0152-0, scully, Ralph Panday, Arvind Elango, Rajula Willis, Nicholas A eng R01 CA095175/CA/NCI NIH HHS/P50 CA168504/CA/NCI NIH HHS/R01 CA217991/CA/NCI NIH HHS/R01 GM134425/GM/NIGMS NIH HHS/R21 ES027776/ES/NIEHS NIH HHS/F31 CA260794/CA/NCI NIH HHS/Research Support, N.I.H., Extramural Review England 2019/07/03 Nat Rev Mol Cell Biol. 2019 Nov;20(11):698–714. doi: 10.1038/s41580-019-0152-0. Epub 2019 Jul 1.

Sharp DJ, Rogers GC, Scholey JM. Cytoplasmic dynein is required for poleward chromosome movement during mitosis in Drosophila embryos. Nat Cell Biol. 2000; 2(12):922–30. https://www.ncbi.nlm.nih.gov/pubmed/11146657, doi: 10.1038/35046574, sharp, J J Rogers, G C Scholey, J M eng GM19262/GM/NIGMS NIH HHS/GM55507/GM/NIGMS NIH HHS/Research Support, U.S. Gov’t, P.H.S. England 2001/01/09 Nat Cell Biol. 2000 Dec;2(12):922–30. doi: 10.1038/35046574.

Shepherd TR, D. RR, Huang H, Wamhoff EC, Bathe M. Bioproduction of pure, kilobase-scale single-stranded DNA. Sci Rep. 2019; 9(1):6121. https://www.ncbi.nlm.nih.gov/pubmed/30992517, doi: 10.1038/s41598-019-42665-1, shepherd, Tyson R Du, Rebecca R Huang, Hellen Wamhoff, Eike-Christian Bathe, Mark eng P30 ES002109/ES/NIEHS NIH HHS/R01 MH112694/MH/NIMH NIH HHS/R21 EB026008/EB/NIBIB NIH HHS/Research Support, N.I.H., Extramural Research Support, U.S. Gov’t, Non-P.H.S. England 2019/04/18 Sci Rep. 2019 Apr 16;9(1):6121. doi: 10.1038/s41598-019-42665-1.

Shirk BD, Rodriguez CZ, Pacheco MO, McTyer JB, Shirk PD, Stoppel WL. Enhancing CRISPR Homology Directed Repair in IAL-PiD2 Insect Cells via Reagent Delivery Optimization and Cell Synchronization. Biochem Eng J. 2026; 225. https://www.ncbi.nlm.nih.gov/pubmed/41648661, doi: 10.1016/j.bej.2025.109906, shirk, Bryce D Rodriguez, Cecilia Z Pacheco, Marisa O McTyer, Jasmine B Shirk, Paul D Stoppel, Whitney L eng R35 GM147041/GM/NIGMS NIH HHS/Netherlands 2026/02/06 Biochem Eng J. 2026 Jan;225:109906. doi: 10.1016/j.bej.2025.109906. Epub 2025 Aug 26.

Shy BR, Vykunta VS, Ha A, Talbot A, Roth TL, Nguyen DN, Pfeifer WG, Chen YY, Blaeschke F, Shifrut E, Vedova S, Mamedov MR, Chung JJ, Li H, Yu R, Wu D, Wolf J, Martin TG, Castro CE, Ye L, et al. High-yield genome engineering in primary cells using a hybrid ssDNA repair template and small-molecule cocktails. Nat Biotechnol. 2023; 41(4):521–531. https://www.ncbi.nlm.nih.gov/pubmed/36008610, doi: 10.1038/s41587-022-01418-8, shy, Brian R Vykunta, Vivasvan S Ha, Alvin Talbot, Alexis Roth, Theodore L Nguyen, David N Pfeifer, Wolfgang G Chen, Yan Yi Blaeschke, Franziska Shifrut, Eric Vedova, Shane Mamedov, Murad R Chung, Jing-Yi Jing Li, Hong Yu, Ruby Wu, David Wolf, Jeffrey Martin, Thomas G Castro, Carlos E Ye, Lumeng Esensten, Jonathan H Eyquem, Justin Marson, Alexander eng F30 DK120213/DK/NIDDK NIH HHS/K08 AI153767/AI/NIAID NIH HHS/T32 GM007618/GM/NIGMS NIH HHS/L40 AI140341/AI/NIAID NIH HHS/L30 TR002983/TR/NCATS NIH HHS/S10 OD025096/OD/NIH HHS/S10 RR028962/RR/NCRR NIH HHS/P01 AI155393/AI/NIAID NIH HHS/L30 AI140341/AI/NIAID NIH HHS/P01 AI138962/AI/NIAID NIH HHS/T32 AI007334/AI/NIAID NIH HHS/T32 DK007418/DK/NIDDK NIH HHS/Research Support, N.I.H., Extramural Research Support, Non-U.S. Gov’t 2022/08/26 Nat Biotechnol. 2023 Apr;41(4):521–531. doi: 10.1038/s41587-022-01418-8. Epub 2022 Aug 25.

Sievers QL, Petzold G, Bunker RD, Renneville A, Slabicki M, Liddicoat BJ, Abdulrahman W, Mikkelsen T, Ebert BL, Thoma NH. Defining the human C2H2 zinc finger degrome targeted by thalidomide analogs through CRBN. Science. 2018; 362(6414). https://www.ncbi.nlm.nih.gov/pubmed/30385546, doi: 10.1126/science.aat0572, sievers, Quinlan L Petzold, Georg Bunker, Richard D Renneville, Aline Slabicki, Mikolaj Liddicoat, Brian J Abdulrahman, Wassim Mikkelsen, Tarjei Ebert, Benjamin L Thoma, Nicolas H eng P50 CA206963/CA/NCI NIH HHS/R01 HL082945/HL/NHLBI NIH HHS/HHMI/Howard Hughes Medical Institute/T32 GM007753/GM/NIGMS NIH HHS/P01 CA066996/CA/NCI NIH HHS/P01 CA108631/CA/NCI NIH HHS/R01 HL082941/HL/NHLBI NIH HHS/Research Support, N.I.H., Extramural Research Support, Non-U.S. Gov’t 2018/11/06 Science. 2018 Nov 2;362(6414):eaat0572. doi: 10.1126/science.aat0572.

Song J, Yang D, Xu J, Zhu T, Chen YE, Zhang J. RS-1 enhances CRISPR/Cas9- and TALEN-mediated knock-in efficiency. Nat Commun. 2016; 7:10548. https://www.ncbi.nlm.nih.gov/pubmed/26817820, doi: 10.1038/ncomms10548, song, Jun Yang, Dongshan Xu, Jie Zhu, Tianqing Chen, Y Eugene Zhang, Jifeng eng R01 HL117491/HL/NHLBI NIH HHS/R01 HL129778/HL/NHLBI NIH HHS/R01HL117491/HL/NHLBI NIH HHS/R01HL129778/HL/NHLBI NIH HHS/Research Support, N.I.H., Extramural England 2016/01/29 Nat Commun. 2016 Jan 28;7:10548. doi: 10.1038/ncomms10548.

Takata M, Sasaki MS, Sonoda E, Morrison C, Hashimoto M, Utsumi H, Yamaguchi-Iwai Y, Shinohara A, Takeda S. Homologous recombination and non-homologous end-joining pathways of DNA double-strand break repair have overlapping roles in the maintenance of chromosomal integrity in vertebrate cells. EMBO J. 1998; 17(18):5497–508. https://www.ncbi.nlm.nih.gov/pubmed/9736627, doi: 10.1093/emboj/17.18.5497, takata, M Sasaki, S S Sonoda, E Morrison, C Hashimoto, M Utsumi, H Yamaguchi-Iwai, Y Shinohara, A Takeda, S eng Research Support, Non-U.S. Gov’t England 1998/09/16 EMBO J. 1998 Sep 15;17(18):5497–508. doi: 10.1093/emboj/17.18.5497.

Taylor GCA, Macdonald L, Szulc NA, Gudauskaite E, Hernandez Moran B, Brisbane JM, Donald M, Taylor E, Zheng D, Gu B, Mill P, Yeyati PL, Pokrzywa W, Ribeiro de Almeida C, Wood AJ. Tissue-specific consequences of tag fusions on protein expression in transgenic mice. PLoS Genet. 2025; 21(8):e1011830. https://www.ncbi.nlm.nih.gov/pubmed/40853976, doi: 10.1371/journal.pgen.1011830, taylor, GillianA A Macdonald, Lewis Szulc, Natalia A Gudauskaite, Evelina Hernandez Moran, Brianda Brisbane, Jennifer M Donald, Molly Taylor, Ella Zheng, Dejin Gu, Bin Mill, Pleasantine Yeyati, Patricia L Pokrzywa, Wojciech Ribeiro de Almeida, Claudia Wood, Andrew J eng WT_/Wellcome Trust/United Kingdom R03 CA286646/CA/NCI NIH HHS/R37 CA269076/CA/NCI NIH HHS/2025/08/25 20:45 PLoS Genet. 2025 Aug 25;21(8):e1011830. doi: 10.1371/journal.pgen.1011830. eCollection 2025 Aug.

Tovell H, Testa A, Maniaci C, Zhou H, Prescott AR, Macartney T, Ciulli A, Alessi DR. Rapid and Reversible Knockdown of Endogenously Tagged Endosomal Proteins via an Optimized HaloPROTAC Degrader. ACS Chem Biol. 2019; 14(5):882–892. https://www.ncbi.nlm.nih.gov/pubmed/30978004, doi: 10.1021/acschembio.8b01016, tovell, Hannah Testa, Andrea Maniaci, Chiara Zhou, Houjiang Prescott, Alan R Macartney, Thomas Ciulli, Alessio Alessi, Dario R eng MC_U127015387/MRC_/Medical Research Council/United Kingdom MC_UU_00018/1/MRC_/Medical Research Council/United Kingdom MC_UU_12016/2/MRC_/Medical Research Council/United Kingdom BB_/Biotechnology and Biological Sciences Research Council/United Kingdom Research Support, Non-U.S. Gov’t 2019/04/13 ACS Chem Biol. 2019 May 17;14(5):882–892. doi: 10.1021/acschembio.8b01016. Epub 2019 Apr 22.

Vaisberg EA, Koonce MP, McIntosh JR. Cytoplasmic dynein plays a role in mammalian mitotic spindle formation. J Cell Biol. 1993; 123(4):849–58. https://www.ncbi.nlm.nih.gov/pubmed/8227145, doi: 10.1083/jcb.123.4.849, vaisberg, A A Koonce, P P McIntosh, J R eng GM 36663/GM/NIGMS NIH HHS/Research Support, U.S. Gov’t, P.H.S. 1993/11/01 J Cell Biol. 1993 Nov;123(4):849–58. doi: 10.1083/jcb.123.4.849.

Velangani HG, Ghosh A, Singh S, Kiran S. Strategies and considerations for the generation of ssDNA-Based HDR templates for CRISPR-based genome editing. BMC Genomics. 2026; 27(1). https://www.ncbi.nlm.nih.gov/pubmed/41634544, doi: 10.1186/s12864-025-12406-y, velangani, Harshitha Golagana Ghosh, Aditi Singh, Sudiksha Kiran, Shashi eng [EMDR/SG/15/2024-01-03223 (E-File No. 222379)]/Indian Council of Medical Research/FILE NO. CRG/2023/007695/ANRF, India/Review England 2026/02/04 BMC Genomics. 2026 Feb 3;27(1):245. doi: 10.1186/s12864-025-12406-y.

Westermann L, Li Y, Gocmen B, Niedermoser M, Rhein K, Jahn J, Cascante I, Scholer F, Moser N, Neubauer B, Hofherr A, Behrens YL, Gohring G, Kottgen A, Kottgen M, Busch T. Wildtype heterogeneity contributes to clonal variability in genome edited cells. Sci Rep. 2022; 12(1):18211. https://www.ncbi.nlm.nih.gov/pubmed/36307508, doi: 10.1038/s41598-022-22885-8, westermann, Lukas Li, Yong Gocmen, Burulca Niedermoser, Matthias Rhein, Kilian Jahn, Johannes Cascante, Isabel Scholer, Felix Moser, Niklas Neubauer, Bjorn Hofherr, Alexis Behrens, Yvonne Lisa Gohring, Gudrun Kottgen, Anna Kottgen, Michael Busch, Tilman eng NakSys/Else Kroner-Fresenius-Stiftung/SFB1453/Deutsche Forschungsgemeinschaft/TRR152/Deutsche Forschungsgemeinschaft/CIBSS/Deutsche Forschungsgemeinschaft/Research Support, Non-U.S. Gov’t England 2022/10/29 Sci Rep. 2022 Oct 28;12(1):18211. doi: 10.1038/s41598-022-22885-8.

Wienert B, Nguyen DN, Guenther A, Feng SJ, Locke MN, Wyman SK, Shin J, Kazane KR, Gregory GL, Carter MAM, Wright F, Conklin BR, Marson A, Richardson CD, Corn JE. Timed inhibition of CDC7 increases CRISPR-Cas9 mediated templated repair. Nat Commun. 2020; 11(1):2109. https://www.ncbi.nlm.nih.gov/pubmed/32355159, doi: 10.1038/s41467-020-15845-1, wienert, Beeke Nguyen, David N Guenther, Alexis Feng, Sharon J Locke, Melissa N Wyman, Stacia K Shin, Jiyung Kazane, Katelynn R Gregory, Georgia L Carter, MatthewM M Wright, Francis Conklin, Bruce R Marson, Alex Richardson, Chris D Corn, Jacob E eng DP2 HL141006/HL/NHLBI NIH HHS/L40 AI140341/AI/NIAID NIH HHS/P30 AI027763/AI/NIAID NIH HHS/Research Support, N.I.H., Extramural Research Support, Non-U.S. Gov’t England 2020/05/02 Nat Commun. 2020 Apr 30;11(1):2109. doi: 10.1038/s41467-020-15845-1.

Xie K, Starzyk J, Majumdar I, Wang J, Rincones K, Tran T, Lee D, Niemi S, Famiglietti J, Suter B, Shan R, Wu H. Efficient non-viral immune cell engineering using circular single-stranded DNA-mediated genomic integration. Nat Biotechnol. 2025; 43(11):1821–1832. https://www.ncbi.nlm.nih.gov/pubmed/39663372, doi: 10.1038/s41587-024-02504-9, xie, Keqiang Starzyk, Jakob Majumdar, Ishita Wang, Jiao Rincones, Katerina Tran, Thao Lee, Danna Niemi, Sarah Famiglietti, John Suter, Bernhard Shan, Richard Wu, Hao eng 2024/12/12 Nat Biotechnol. 2025 Nov;43(11):1821–1832. doi: 10.1038/s41587-024-02504-9. Epub 2024 Dec 11.

Xu J, Liang Y, Li N, Dang S, Jiang A, Liu Y, Guo Y, Yang X, Yuan Y, Zhang X, Yang Y, Du Y, Shi A, Liu X, Li D, He K. Clathrin-associated carriers enable recycling through a kiss-and-run mechanism. Nat Cell Biol. 2024; 26(10):1652–1668. https://www.ncbi.nlm.nih.gov/pubmed/39300312, doi: 10.1038/s41556-024-01499-4, xu, Jiachao Liang, Yu Li, Nan Dang, Song Jiang, Amin Liu, Yiqun Guo, Yuting Yang, Xiaoyu Yuan, Yi Zhang, Xinyi Yang, Yaran Du, Yongtao Shi, Anbing Liu, Xiaoyun Li, Dong He, Kangmin eng 32321004, 32321004/National Natural Science Foundation of China (National Science Foundation of China)/England 2024/09/20 Nat Cell Biol. 2024 Oct;26(10):1652–1668. doi: 10.1038/s41556-024-01499-4. Epub 2024 Sep 19.

Yadav S, Puthenveedu MA, Linstedt AD. Golgin160 recruits the dynein motor to position the Golgi apparatus. Dev Cell. 2012; 23(1):153–65. https://www.ncbi.nlm.nih.gov/pubmed/22814606, doi: 10.1016/j.devcel.2012.05.023, yadav, Smita Puthenveedu, Manojkumar A Linstedt, Adam D eng R00 DA024698/DA/NIDA NIH HHS/R01 GM056779/GM/NIGMS NIH HHS/GM056779/GM/NIGMS NIH HHS/DA024698/DA/NIDA NIH HHS/Research Support, N.I.H., Extramural 2012/07/21 Dev Cell. 2012 Jul 17;23(1):153–65. doi: 10.1016/j.devcel.2012.05.023.

Yang D, Scavuzzo MA, Chmielowiec J, Sharp R, Bajic A, Borowiak M. Enrichment of G2/M cell cycle phase in human pluripotent stem cells enhances HDR-mediated gene repair with customizable endonucleases. Sci Rep. 2016; 6:21264. https://www.ncbi.nlm.nih.gov/pubmed/26887909, doi: 10.1038/srep21264, yang, Diane Scavuzzo, Marissa A Chmielowiec, Jolanta Sharp, Robert Bajic, Aleksandar Borowiak, Malgorzata eng P30-DK079638/DK/NIDDK NIH HHS/P30 CA125123/CA/NCI NIH HHS/P30 DK079638/DK/NIDDK NIH HHS/P30 AI036211/AI/NIAID NIH HHS/S10 RR024574/RR/NCRR NIH HHS/Research Support, N.I.H., Extramural Research Support, Non-U.S. Gov’t England 2016/02/19 Sci Rep. 2016 Feb 18;6:21264. doi: 10.1038/srep21264.

Yesbolatova A, Saito Y, Kitamoto N, Makino-Itou H, Ajima R, Nakano R, Nakaoka H, Fukui K, Gamo K, Tominari Y, Takeuchi H, Saga Y, Hayashi KI, Kanemaki MT. The auxin-inducible degron 2 technology provides sharp degradation control in yeast, mammalian cells, and mice. Nat Commun. 2020; 11(1):5701. https://www.ncbi.nlm.nih.gov/pubmed/33177522, doi: 10.1038/s41467-020-19532-z, yesbolatova, Aisha Saito, Yuichiro Kitamoto, Naomi Makino-Itou, Hatsune Ajima, Rieko Nakano, Risako Nakaoka, Hirofumi Fukui, Kosuke Gamo, Kanae Tominari, Yusuke Takeuchi, Haruki Saga, Yumiko Hayashi, Ken-Ichiro Kanemaki, Masato T eng Research Support, Non-U.S. Gov’t England 2020/11/13 Nat Commun. 2020 Nov 11;11(1):5701. doi: 10.1038/s41467-020-19532-z.

Zhang JP, Li XL, Li GH, Chen W, Arakaki C, Botimer GD, Baylink D, Zhang L, Wen W, Fu YW, Xu J, Chun N, Yuan W, Cheng T, Zhang XB. Efficient precise knockin with a double cut HDR donor after CRISPR/Cas9-mediated double-stranded DNA cleavage. Genome Biol. 2017; 18(1):35. https://www.ncbi.nlm.nih.gov/pubmed/28219395, doi: 10.1186/s13059-017-1164-8, zhang, Jian-Ping Li, Xiao-Lan Li, Guo-Hua Chen, Wanqiu Arakaki, Cameron Botimer, Gary D Baylink, David Zhang, Lu Wen, Wei Fu, Ya-Wen Xu, Jing Chun, Noah Yuan, Weiping Cheng, Tao Zhang, Xiao-Bing eng England 2017/02/22 Genome Biol. 2017 Feb 20;18(1):35. doi: 10.1186/s13059-017-1164-8.

Zhang W, Chen Y, Yang J, Zhang J, Yu J, Wang M, Zhao X, Wei K, Wan X, Xu X, Jiang Y, Chen J, Gao S, Mao Z. A high-throughput small molecule screen identifies farrerol as a potentiator of CRISPR/Cas9-mediated genome editing. Elife. 2020; 9. https://www.ncbi.nlm.nih.gov/pubmed/32644042, doi: 10.7554/eLife.56008, zhang, Weina Chen, Yu Yang, Jiaqing Zhang, Jing Yu, Jiayu Wang, Mengting Zhao, Xiaodong Wei, Ke Wan, Xiaoping Xu, Xiaojun Jiang, Ying Chen, Jiayu Gao, Shaorong Mao, Zhiyong eng 2018YFC2000100/Chinese National Program on Key Basic Research Project/International 19JC1415300/the key project of the Science and Technology of Shanghai Municipality/International 19QA1409600/the Shanghai Rising-Star Program/International 2017ZZ02015/the Shanghai Municipal Medical and Health Discipline Construction Projects/International 2018QNRC001/the Young Elite Scientist Sponsorship Program by CAST/International 2017YFA010330/Chinese National Program on Key Basic Research Project/International 2016YFA0100400/Chinese National Program on Key Basic Research Project/International 31871438/the National Science Foundation of China/International 81972457/the National Science Foundation of China/International 31721003/the National Science Foundation of China/International 31871446/the National Science Foundation of China/International 19XD1403000/the Fundamental Research Funds for the Central Universities, Program of Shanghai Academic Research Leader/International 19SG18/Shuguang Program” of Shanghai Education Development Foundation and Shanghai Municipal Education Commission”/International 2017YFA0103300/Chinese National Program on Key Basic Research Project/International 31871438/National Science Foundation of China/International 81972457/National Science Foundation of China/International 31721003/National Science Foundation of China/International 31871446/National Science Foundation of China/International 19XD1403000/Program of Shanghai Academic Research Leader/International 19SG18/”Shuguang Program” of Shanghai Education Development Foundation and Shanghai Municipal Education Commission/International 19QA1409600/Shanghai Rising-Star Program/International 2017ZZ02015/Shanghai Municipal Medical and Health Discipline Construction Projects/International Research Support, Non-U.S. Gov’t England 2020/07/10 Elife. 2020 Jul 9;9:e56008. doi: 10.7554/eLife.56008.

Zimmerman SP, DeGraw LB, Counter CM. The essential clathrin adapter protein complex-2 is tumor suppressive specifically in vivo. Nat Commun. 2025; 16(1):2254. https://www.ncbi.nlm.nih.gov/pubmed/40050266, doi: 10.1038/s41467-025-57521-2, zimmerman, Seth P DeGraw, Lili B Counter, Christopher M eng F32CA236183/U.S. Department of Health & Human Services | NIH | National Cancer Institute (NCI)/F32 CA236183/CA/NCI NIH HHS/R01 CA269272/CA/NCI NIH HHS/T32 CA009111/CA/NCI NIH HHS/P30 CA014236/CA/NCI NIH HHS/P30CA014236/U.S. Department of Health & Human Services | NIH | National Cancer Institute (NCI)/R01CA269272/U.S. Department of Health & Human Services | NIH | National Cancer Institute (NCI)/England 2025/03/07 Nat Commun. 2025 Mar 6;16(1):2254. doi: 10.1038/s41467-025-57521-2.

